# E-cadherin is sorted by Rab7 and Snx16 for polarised secretion via Myosin V

**DOI:** 10.1101/2022.02.17.480827

**Authors:** Dajana Tanasic, Nicola Berns, Veit Riechmann

## Abstract

E-cadherin has a fundamental role in epithelial tissues by providing cell-cell adhesion. Epithelial homeostasis relies on polarised E-cadherin exocytosis to the lateral plasma membrane, however the secretion mechanisms are unknown. Epithelial plasticity depends on constant E-cadherin endocytosis and recycling, but it is unclear how recycling is facilitated. Here we use the *Drosophila* follicular epithelium to analyse E-cadherin recycling and secretion. We identify endosomes in the apical region of the epithelium, in which newly translated and endocytosed E-cadherin converge for polarised E-cadherin secretion. Our data provide evidence that Rab7 recruits Snx16 to these endosomes, and that Snx16 moves E-cadherin via tubulation into the Rab11 compartment. Rab11 forms E-cadherin transport vesicles by recruiting its effector Myosin V. We show in living follicles how Myosin V transports E-cadherin along an apical actin network to the zonula adherence. An additional secretion pathway exists in the basal epithelium, where Myosin V moves E-cadherin vesicles along parallel actin bundles to the plasma membrane.

## Introduction

The integral membrane protein E-cadherin provides homophilic cell-cell adhesion in epithelial cells by forming contact sites called adherens junctions. E-cadherin assembles two types of adherens junctions: punctate adherens junctions which distribute along the lateral plasma membrane (PM) and belt-like adherens junctions. The latter are established in the apical-most region of the lateral membrane, where they form the zonula adherens (ZA). In both types of junctions E-cadherin links the actin cytoskeleton to the PM via its association with *β*- and *α*- catenin (Harris and Tepass, 2010; Takeichi, 2014).

Epithelial tissues require a high degree of plasticity because they are continuously renewed and exposed to mechanical stress. Plasticity relies on the constant exchange of E-cadherin molecules at the PM, which is achieved by trafficking in membrane vesicles. Many morphogenetic processes rely on changes in the E-cadherin trafficking system. For instance, during epithelial-mesenchymal transitions (EMT), which are a driving force for embryonic development, E-cadherin distribution at the PM changes to allow cell migration. Defects in E-cadherin trafficking are the cause of many diseases including cancer, where the loss of E-cadherin mediated cell-cell adhesion is a critical step during metastasis (Dongre and Weinberg, 2019; Thiery et al., 2009).

Studies in mammalian cells have shown that E-cadherin is constantly endocytosed and recycled (de Beco et al., 2009; Le et al., 1999). After endocytosis, vesicles fuse with the early endosome, which is organised by the small GTPase Rab5 (Brüser and Bogdan, 2017; Cadwell et al., 2016). Within endosomes, E-cadherin is either sorted for degradation in lysosomes or for recycling back to the PM (Brüser and Bogdan, 2017). Endosomes harbour multiple protein complexes that cooperate in cargo recognition and distribution into various pathways. Cargos that are targeted for degradation are sent to the endolysosomal pathway. In *Drosophila*, RabX1 was recently shown to promote the degradation of endocytosed proteins by establishing highly efficient endolysosomes (Laiouar et al., 2020). Cargos destined for recycling are recognised by retrieval complexes, which are called retromer, retriever, CCC and ESCAPE-1. These complexes sort cargos into the different endosomal recycling branches (Cullen and Steinberg, 2018; Weeratunga et al., 2020). At the end of these branches tubulo- vesicular carriers are generated, in which cargos are transported to the Golgi or to the PM (Simonetti and Cullen, 2019). It is not known how E-cadherin is retrieved from degradation but several good candidates have been identified. These include the F-Bar proteins Cip4 and Nostrin (Zobel et al., 2015) and the sorting nexins Snx1 (Bryant et al., 2007), Snx4 (Solis et al., 2013) and Snx16 (Wang et al., 2019; Xu et al., 2017). Various studies indicate that endosomal sorting of E-cadherin includes passage through a Rab11 compartment (Bogard et al., 2007; Classen et al., 2005; Desclozeaux et al., 2008; Le Droguen et al., 2015; Lock and Stow, 2005; Pirraglia et al., 2010; Roeth et al., 2009). The strong intracellular accumulation in *Rab11* mutant cells suggests that this passage is central for E-cadherin secretion (Woichansky et al., 2016; Xu et al., 2011).

Live imaging in mammalian cells indicated that the Rab11 compartment receives also newly synthesised E-cadherin (Desclozeaux et al., 2008; Lock and Stow, 2005). This suggests that the endosomal sorting machinery is used to target new E-cadherin to the PM. Consistent with this, we previously reported that the intracellular E-cadherin aggregates in *Rab11* mutant cells consist to a large extent of newly translated protein (Woichansky et al., 2016). The Rab11 compartment is therefore central for the secretion of both new and recycled E-cadherin. However, it is unclear how E-cadherin traffics into the Rab11 compartment.

It is also unknown how E-cadherin vesicles are transported from the Rab11 compartment to the PM. One critical Rab11 effector is Sec15 (Wu et al., 2005; Zhang et al., 2004), a component of the exocyst complex, which facilitates vesicle fusion with the PM (Grindstaff et al., 1998; Yeaman et al., 2004). The link between E-cadherin and the exocyst complex is thought to be provided by *β*-catenin, which binds directly to E-cadherin and to the exocyst component Sec10 (Langevin et al., 2005). We proposed two pathways along which E-cadherin transport to the PM may occur: “apicolateral exocytosis”, which delivers E-cadherin to the ZA and “lateral exocytosis”, which provides the entire lateral membrane with E-cadherin for punctate adherence junctions formation (Woichansky et al., 2016). Within the lateral membrane, E-cadherin is distributed by an apically directed “cadherin flow” (Kametani and Takeichi, 2007; Woichansky et al., 2016). While the membrane flow has been shown to be actin-dependent, it is unclear on what kind of cytoskeletal tracks DE-cadherin is transported to the PM.

In this study, we analyse DE-cadherin trafficking upstream and downstream of Rab11. We identify apical endosomes, in which the pathways for DE-cadherin recycling and synthesis intersect. Our data suggest that Rab7 recruits Snx16 to these endosomes, and that Snx16 moves DE-cadherin into the Rab11 compartment. Here, Rab11 recruits Myosin V (MyoV) to facilitate DE-cadherin exocytosis along an apical F-actin network to the ZA. A second exocytosis route exists in the basal-most region of the epithelium, where MyoV transports DE-cadherin along parallel F-actin arrays.

## Results

### Endocytosed and newly translated DE-cadherin converge in an apical Rab11 compartment

We use the epithelium of the *Drosophila* ovary to analyse the trafficking of E-cadherin, which is called DE-cadherin in the fly. The follicular epithelium is a monolayer surrounding the cells of the germline. Its apical membrane faces the germline cells, and its ZA marks the border between the apical and lateral membrane domain. Ovaries harbour follicles of different developmental stages. Here we focus on stage 8 and 9 follicles, in which the epithelial cells have ceased cell proliferation but are not yet differentiated (Horne-Badovinac and Bilder, 2005). Fig. S1 explains how we analysed protein distributions in follicles. Optical confocal sections were taken sagittally to show the epithelium along its apico-basal axis and perpendicular to the apico-basal axis showing the epithelium in apical or basal views.

To localise where in the follicular epithelium DE-cadherin is sorted for recycling we performed pulse-chase endocytosis experiments. We briefly incubated living ovaries with an antibody that binds to the extracellular part of DE-cadherin (Oda et al., 1994). Since the PM was not permeabilised, the antibody bound exclusively to DE-cadherin at the PM. After washing, ovaries were incubated for different periods of time, fixed and stained. After 20 minutes we detected prominent spots of endocytosed (PM-labelled) DE-cadherin (arrows in Fig. 1A,E), which localised close to a previously described network of apical F-actin (Wang and Riechmann, 2007). Interestingly, these spots were surrounded by F-actin suggesting a role of actin in DE-cadherin trafficking. To test whether the DE-cadherin spots represent endosomes we examined how they relate to Rab11, which is central for DE-cadherin recycling. To this end, we expressed an HA-tagged Rab11 protein, which is fully active as revealed by a complete rescue of *Rab11* mutant cell clones (Fig. S2B). HA-Rab11 formed a heterogeneous apical compartment with which PM-labelled DE-cadherin partially overlapped supporting the idea that the PM-labelled DE-cadherin spots represent endosomes (Fig. 1B).

**Figure 1:**
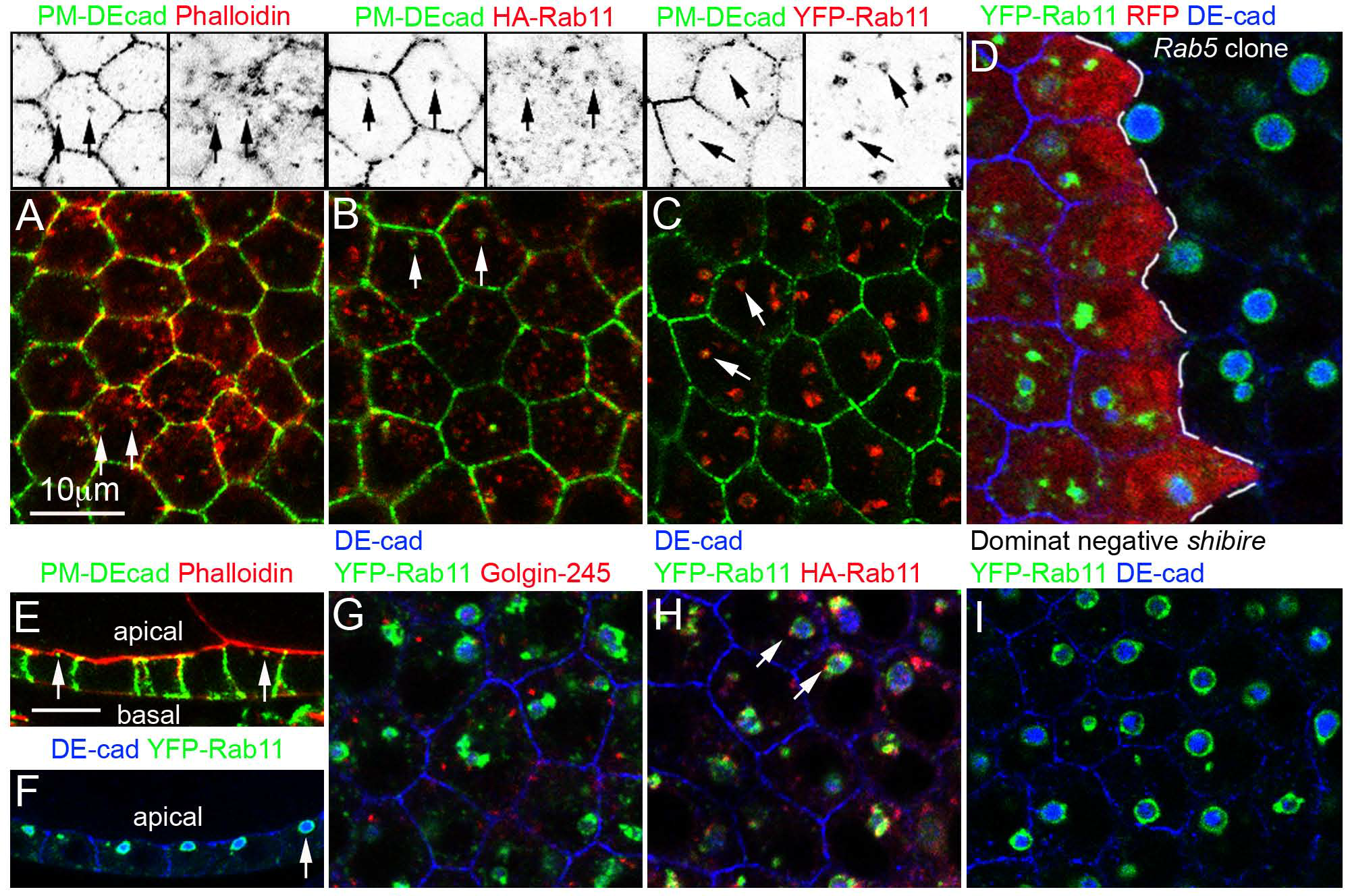
Endocytosed and newly translated DE-cadherin converge in an apical Rab11 compartment. Optical confocal sections through the follicular epithelium. (**A-C**) The apical area of the epithelium is shown. Pulse-chase DE-cadherin endocytosis experiments. Endocytosed (PM-labelled) DE-cadherin is shown in green. The black and white pictures show the individual green and red channels of the pictures below alone. Arrows point at spots where endocytosed DE-cadherin concentrates. (**A**) Wild type follicle that was also stained for Phalloidin (red) to reveal the actin cytoskeleton. 96.67% (± 0.20) of endocytosed DE-cadherin spots localises closely to F-actin (n=176 cells from 4 follicles). (**B**) Follicle expressing HA- Rab11 (red). 81.84% (± 3.11) of the endocytosed DE-cadherin spots overlap with HA-Rab11 (n=159 cells from 5 follicles). (**C**) Follicle expressing YFP-Rab11 (red). Endocytosed DE-cadherin was detected in in 72% (± 2.62) of the YFP-Rab11 vacuoles (n=178 cells from 5 follicles). (**D**) YFP-Rab11 expressing epithelium harbouring a *Rab5* mutant cell clones. Mutant cells are marked by the absence of RFP (red). The clone border is highlighted by the dashed line. *Rab5* mutant cells form YFP-Rab11 vacuoles (green) that are filled with DE-cad (blue) similar to the neighbouring wild type cells. This phenotype was observed in all analysed clones (n=12). (**E-F**) Sagittal section showing the epithelium along its apical-basal axis. (**E**) Pulse-chase endocytosis experiment. Endocytosed DE-cadherin (green) is visible in the apical region of the epithelium (arrows). (**F**) YFP-Rab11 (green) expressing epithelium showing the apical localisation of the DE-cadherin (blue) filled vacuoles (arrow). (**G**) Apical view of a YFP- Rab11 expressing epithelium. DE-cadherin (blue) accumulates within the YFP-Rab11 vacuoles (green). The *trans*-Golgi marker Golgin-245 (red) shows almost no overlap with YFP- Rab11. (**H**) Follicle co-expressing YFP-Rab11 (green) and HA-Rab11 (red). Arrows point at HA-Rab11 spots that are visible at the edge of the YFP-Rab11 vacuoles. (**I**) Follicle co-expressing YFP-Rab11 (green) and a dominant-negative form of *shibire*. DE-cadherin (blue) was detected in 85.95% (± 3.73) of YFP-Rab11 vacuoles (n=207 cells from 6 follicles).

We also included a YFP-tagged Rab11 protein in our analysis which, surprisingly, led to the formation of large vacuoles (Fig. 1G). These vacuoles were confined by YFP-Rab11 staining and showed a strong DE-cadherin signal inside. This suggests that DE-cadherin trafficking is blocked within the YFP-Rab11 compartment. We found no gross changes in cell shape and ZA formation, which suggests that the endogenous Rab11 supports sufficient DE-cadherin trafficking to the PM. To analyse how a functional Rab11 protein contributes to trafficking in YFP-Rab11 follicles we co-expressed HA-Rab11. HA-Rab11 localised in a punctate pattern at the edge of the YFP-Rab11 vacuoles (arrows in Fig. 1H). This raises the possibility that HA-Rab11 facilitates DE-cadherin transport from the vacuoles to the PM. The vacuoles might thus not represent a dead end in DE-cadherin transport, but rather a blockage that is released at their periphery by active Rab11. Since these vacuoles block trafficking in a well-defined and enlarged compartment, we used them to analyse DE-cadherin trafficking within the Rab11 compartment.

The YFP-Rab11 vacuoles localise to the apical area of the epithelium (Fig. 1F). In this region we also detected the prominent spots of endocytosed DE-cadherin (Fig. 1E). To test if the vacuoles are integrated into the recycling pathway we performed endocytosis assays, and detected DE-cadherin after 30 minutes within the vacuoles (Fig. 1C). Rab11 is not only required for recycling of endocytosed DE-cadherin but also for the secretion of new protein. To test if newly synthesised DE-cadherin also accumulates in the vacuoles we blocked the import of endocytosed DE-cadherin into the YFP-Rab11 vacuoles by removing Rab5, which is required for the fusion of endocytic vesicles with the endosome. We generated genetic mosaic epithelia harbouring homozygous *Rab5* mutant cells. To exclude residual Rab5 function we used a null allele (Wucherpfennig et al., 2003). To avoid Rab5 perdurance we focused on clones covering at least one third of the epithelium, which implies that they have undergone eight cell divisions after clone induction (Margolis and Spradling, 1995). Notably, the YFP- Rab11 vacuoles in *Rab5* mutant cell clones still accumulated DE-cadherin indicating that the YFP-Rab11 vacuoles harbour also newly translated protein (Fig. 1D). We also interfered with endocytosis by expressing a dominant-negative form of *shibire*, which encodes *Drosophila* Dynamin, a critical protein for scission of clathrin-coated vesicles (Hill et al., 2001). This confirmed that DE-cadherin still accumulates in the YFP-Rab11 vacuoles when its internalisation from the PM is blocked (Fig. 1I). In summary, these data indicate that the apical YFP-Rab11 vacuoles are supplied with endocytosed as well as with newly synthesised DE-cadherin. This suggests that the recycling and biosynthetic pathways for DE-cadherin merge in the YFP-Rab11 vacuoles. In conclusion, our data suggest the existence of an apical Rab11 compartment in which new and recycled DE-cadherin converge.

### Rab11 is required for DE-cadherin export from the Rab7 compartment

To find out which sorting step is defective in the YFP-Rab11 vacuoles we examined the distribution of the endosomal markers Rab5 and Rab7. Interestingly, both proteins largely overlapped with YFP-Rab11 at the periphery of the vacuoles (Fig. 2A). This suggests that the vacuoles resulted from a fusion of the YFP-Rab11, Rab5 and Rab7 compartments. The presence of Rab5 suggests that the early endosome partially fused with the vacuoles. The presence of Rab7 is more difficult to explain since Rab7 has functions in degradation and recycling (see below).

**Figure 2:**
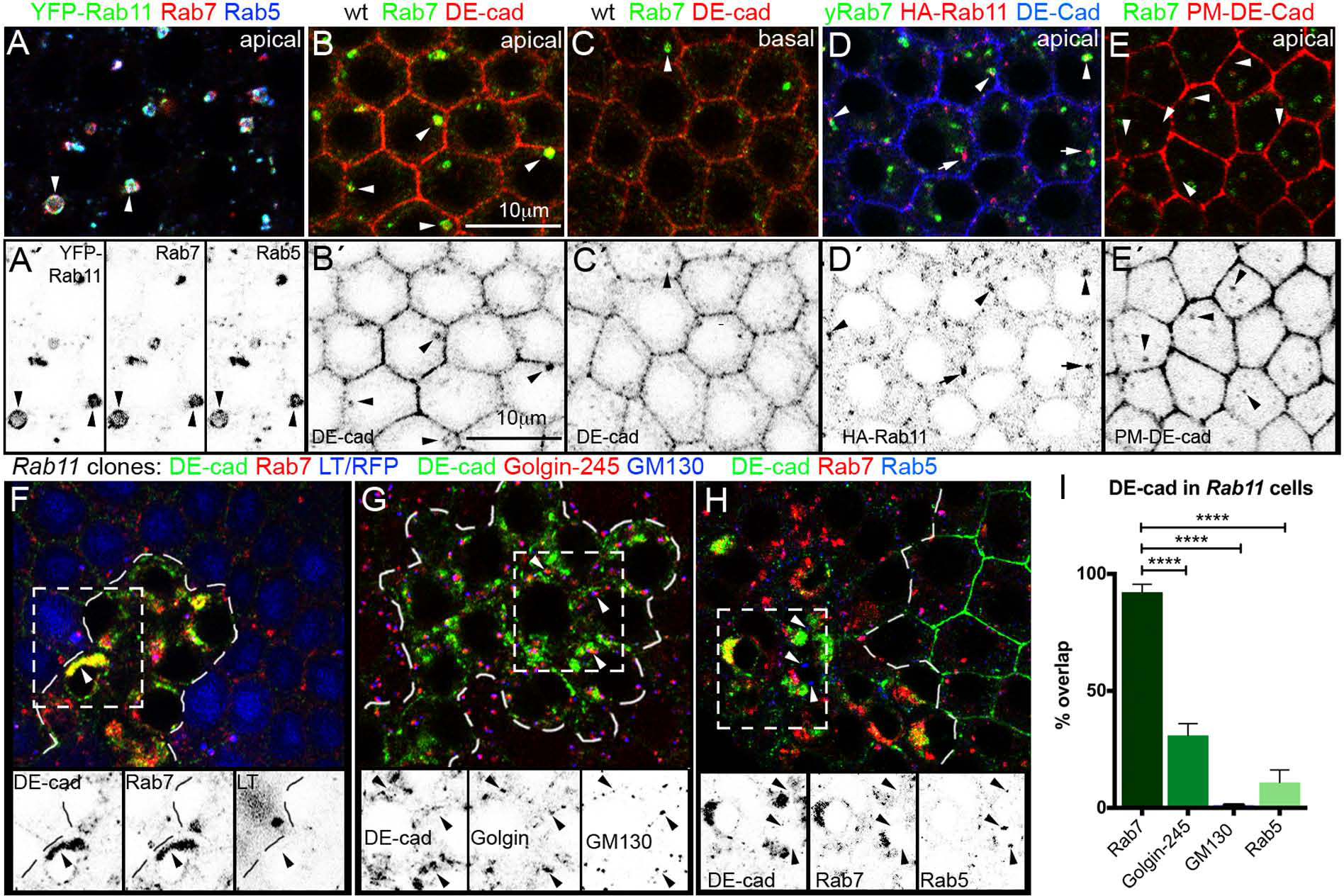
DE-cadherin traffics through endosomal Rab7 and Rab11 compartments. Optical confocal sections through the follicular epithelium. (**A**) Apical confocal section of a follicle expressing YFP-Rab11 (green), stained for the early endosome marker Rab5 (blue) and Rab7 (red). (**Á**) The insets below show single channels. The signals of YFP-Rab11, Rab5 and Rab7 overlap in the vacuoles and in the puncta. (**B**) Apical confocal section of an epithelium stained for Rab7 (green) and DE-cadherin (red). Strong red staining at the PM in the middle shows the DE-cadherin concentration at the ZA. Big Rab7 compartments are detectable in the apical area of the epithelium. DE-cadherin is visible within the Rab7 compartment (arrowheads). (**B’**) DE-cadherin channel alone. (**C**) Basal confocal section of the same epithelium. Fewer big Rab7 compartments are detectable basally. (**Ć**) DE-cadherin channel alone. (**D**) Apical confocal section of an epithelium in which the endogenous Rab7 gene is tagged with GFP (yRab7) and in which HA-Rab11 (red) is expressed. The yRab7 compartments partially overlap with the HA-Rab11 compartments (arrowheads point to the yellow overlap). Overlap with HA-Rab11 was detected in 77% (± 5.74) of the yRab7 compartments (n=172 cells from 6 follicles). HA-Rab11 also forms tubulo-vesicular structures next to the yRab7 compartment (arrow). (**D’**) HA-Rab11 channel alone. (**E**) Pulse-chase endocytosis experiment in which DE-cadherin (red) was detected after 20 minutes within the apical Rab7 compartment (green, arrows). 89.57% (± 2.26) of the DE-cadherin spots co- localised with the Rab7 signal (n=220 cells from 6 follicles). (**É**) DE-cadherin channel alone. (**F-H**) Confocal sections of epithelia harbouring *Rab11* mutant cell clones which are marked by the dotted line. The tissue is stained as indicated. Insets show single channels of the region marked by the dashed box. (**F**) DE-cadherin (green) aggregates in *Rab11* mutant cells show partial and complete overlap with Rab7 (red). However, DE-cadherin aggregates are negative for the lysosomal marker lysotracker (blue), which indicates that they form outside of the degradative part of endosomes. (**G**) DE-cadherin aggregates (green) show almost no overlap with the *cis*-Golgi marker GM130 (blue) and the *trans*-Golgi marker Golgin-245 (red). (**H**) DE-cadherin aggregates in the *Rab11* clone overlap with Rab7 (red) but not with the early endosome marker Rab5 (arrowheads, blue). (**I**) Quantification of the overlap of the DE- cadherin aggregates in *Rab11* mutant cells with Rab7, Golgin-245 (*trans*-Golgi), GM130 (*cis*- Golgi) and Rab5. Data are shown as mean ± SEM. Two-tailed t-test (equal variance, *α* = 0.05) was performed and p values are presented as ****p < 0.0001 (compared groups: Rab7 and DE-cad overlap versus Golgin-245 and DE-cad overlap, Rab7 and DE-cad overlap versus GM130 and DE-cad overlap, Rab7 and DE-cad overlap versus Rab5 and DE-cad overlap, all in *Rab11* clone cells). The quantification was performed with 10 follicles (n=197 clone cells) for Rab7 and DE-cad overlap, 14 follicles (n=276 clone cells) for Golgin-245 and DE-cad overlap, 9 follicles (n=165 clone cells) for GM130 and DE-cad overlap and 7 follicles (n=122 clone cells) for Rab5 and DE-cad overlap.

To examine how Rab7 contributes to DE-cadherin trafficking we first analysed its compartment in wild type epithelia. Rab7 formed a relatively big apical compartment of an average size of 0.88 μm^2^ (± 0.07, n=120 cells from 10 follicles), in which DE-cadherin is detectable (arrowheads in Fig. 2B). Notably, 83% of these big Rab7 compartments localise to the apical and only 18% to the basal region of the cell (Fig. 2B,C). We expressed HA-Rab11 to find out how the apical Rab7 compartment relates to an active Rab11 compartment. HA-Rab11 partially overlapped with Rab7 and was also detected in tubulo-vesicular structures next to it, which suggests a connection of the two compartments (arrows and arrowheads in Fig. 2D). We examined the specificity of the DE-cadherin co-localisation with the yRab7/HA-Rab11 compartment in relation to aPKC protein, which was not expected to associate with this compartment. aPKC co-localisation with yRab7/HA-Rab11 was low indicating that the recruitment of DE-cadherin to this compartment is specific (Fig. S2C-E). To test if the Rab7 compartment is involved in DE-cadherin recycling we performed pulse-chase experiments. Similar to the HA-Rab11 compartment, endocytosed DE-cadherin concentrated after 20 minutes in the Rab7 compartment (Fig. 2E). Taken together, these data indicate that endocytosed DE-cadherin traffics through an apical endosomal system, which includes the compartments of Rab7 and Rab11.

To analyse the functional relationship between the compartments of Rab7 and Rab11 we mapped where DE-cadherin trafficking is blocked when Rab11 is not active. Notably, in cell clones that are mutant for the null allele *Rab11^dFRT^* (Bogard et al., 2007), DE-cadherin aggregates overlapped to a large extent with Rab7 (arrowhead in Fig. 2F). By contrast, markers for Golgi, lysosomes and early endosomes reveal no or only a weak overlap with the DE-cadherin aggregates (Fig. 2F-I). Thus, in the absence of Rab11, DE-cadherin appears to be trapped within the Rab7 compartment. This suggests that Rab11 is required to facilitate the export of DE-cadherin out of the Rab7 compartment.

### Snx16 transports DE-cadherin out of Rab7 endosomes

The finding that DE-cadherin accumulates in *Rab11* mutant cells within the Rab7 compartment raises the question of what functional role Rab7 has for DE-cadherin trafficking. We therefore generated *Rab7* mutant cell clones using a null allele (Cherry et al., 2013). *Rab7* mutant cells showed no severe defects in cell shape and ZA formation, which indicates that DE-cadherin transport to the PM is at least partially possible in the absence of Rab7. However, proper DE-cadherin trafficking is disturbed as revealed by the formation of small DE-cadherin aggregates (arrowheads in Fig. 3A). This indicates a role of Rab7 in DE-cadherin trafficking.

**Figure 3:**
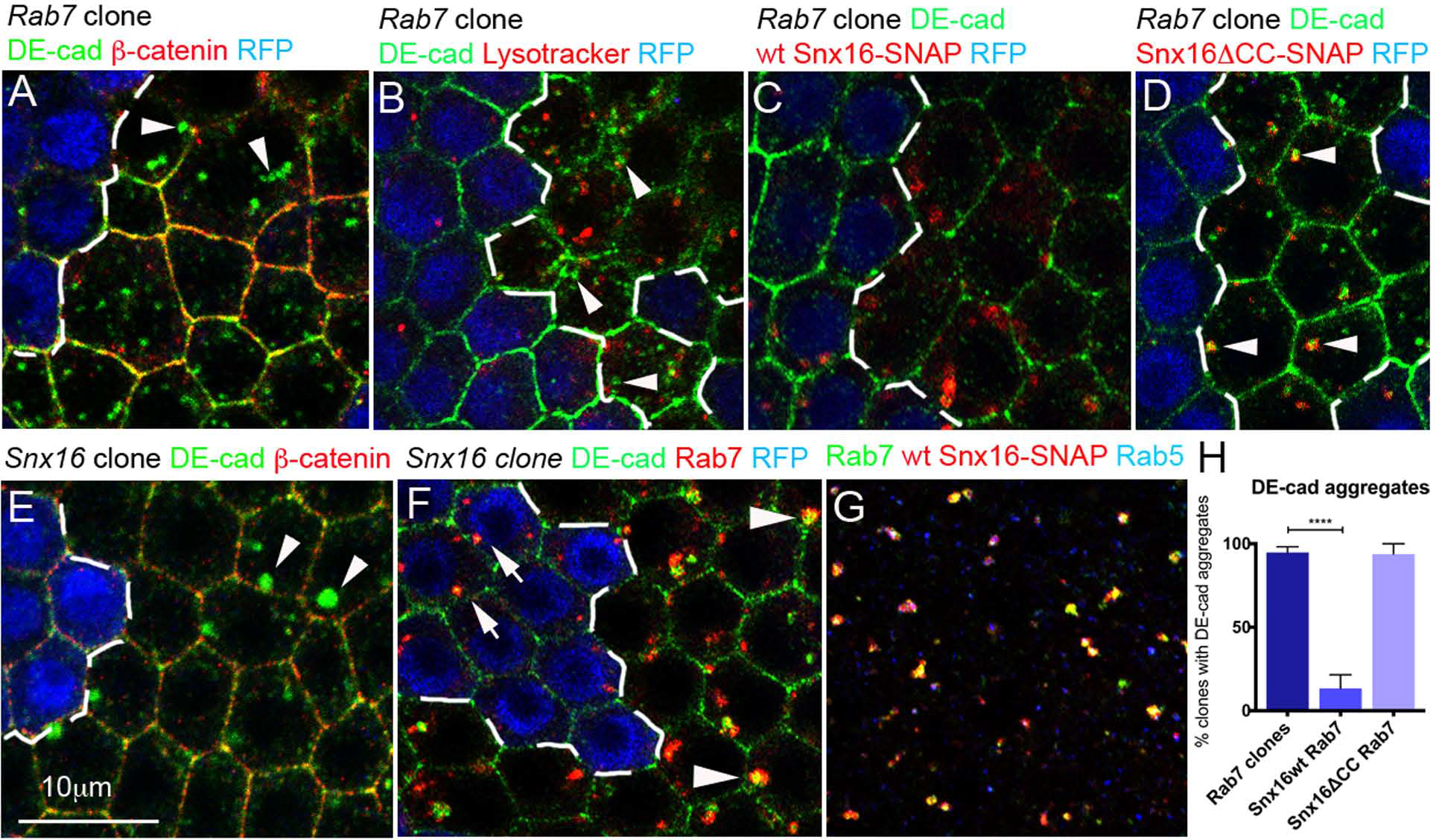
Rab7 and Snx16 control endosomal trafficking. Optical confocal sections through the follicular epithelium. (**A-D**) *Rab7* mutant cell clones, which are marked by the absence of RFP (blue) and whose borders are indicated by the dashed line. (**A**) *Rab7* mutant cells form DE-cadherin aggregates (green). The ZA is formed in *Rab7* mutant cells as revealed by the concentration of DE-cadherin and *β*-catenin (red) at the PM. Aggregates were detected in 95% of 38 analysed cell clones. (**B**) Most DE-cadherin aggregates in *Rab7* mutant cells do not co-localise with the lysosomal marker lysotracker (arrowheads, red). (**C**) Expression of wild type Snx16 (red) in *Rab7* mutant cells rescues DE-cadherin aggregation. (**D**) Expression of an Snx16 protein in which its coiled-coil domain is deleted (Snx16*Δ*CC, red) does not rescue DE-cadherin aggregation in *Rab7* mutant cells. DE-cadherin aggregates form within the mutant Snx16 compartment (arrowheads). This was observed in 87% of the analysed clones (n=15). (**E-F**) *Snx16* mutant cell clones, which are marked by the absence of RFP (blue) and whose borders are indicated by the dashed line (**E**) *Snx16* mutant cell clones stained for *β*- catenin (red) and DE-cadherin (green). DE-cadherin aggregates in the mutant cells. Aggregates were detected in 90% of 34 analysed cell clones. (**F**) *Snx16* mutant cell clones stained for DE-cadherin (green) and Rab7 (red). DE-cadherin is detectable in the Rab7 compartment in wild type (arrows) and *Snx16* mutant cells (arrowheads). In all analysed clones DE-cadherin was detected in the Rab7 compartment (n=23). Note that the DE-cadherin and the Rab7 puncta are bigger in the *Snx16* mutant cells indicating that DE-cadherin trafficking is disturbed within the Rab7 compartment. (**G**) Epithelium expressing wild type Snx16 (red) that was stained for Rab7 (green) and Rab5 (blue). Snx16 signal overlaps almost completely with the Rab7 compartment (88.30 % ± 1.46, n=199 cells from 5 follicles). (**H**) Quantification of DE-cadherin aggregation in *Rab7* mutant cell clones. Quantification of images that were taken in the experiment from which the representative samples are shown in (a), (c) and (d). Data are shown as mean ± SEM. Two-tailed t-test (equal variance, *α* = 0.05) was performed and p values are presented as ****p < 0.0001 (compared groups: *Rab7* clones versus wild type Snx16 expression in *Rab7* clones). The quantification was performed with 30 follicles (n=38 clones) for *Rab7* clones, 17 follicles (n=18 clones) for wild type Snx16 in *Rab7* clones and 8 follicles (n=11 clones) for Snx16*Δ*CC in *Rab7* clones.

Rab7 is known to initiate the endolysosomal pathway and to recruit the retromer complex to endosomes (Rojas et al., 2008; Seaman et al., 2009). The DE-cadherin aggregates in *Rab7* cells could therefore indicate a defect in degradation or recycling. We previously detected an increase in lysotracker puncta in *Rab7* depleted follicle cells, which most likely reflects impaired endolysosomal degradation (Laiouar et al., 2020). Notably, 91% of the DE-cadherin aggregates did not overlap with these lysotracker puncta (n=180 cells from 4 follicles, Fig. 3B). This suggests that most of the DE-cadherin aggregates in *Rab7* cells are not targeted for degradation but rather accumulate because of a defect in a DE-cadherin retrieval.

A good candidate facilitating DE-cadherin retrieval is Snx16 since the protein was shown to bind E-cadherin in endosomes (Xu et al., 2017). Snx16 belongs to the family of sorting nexins, which recognise endosomal cargos and support the formation of membrane tubules (Choi et al., 2004; Hanson and Hong, 2003; Wang et al., 2019). To examine Snx16 localisation we expressed a SNAP-tagged form in the follicular epithelium. Snx16 overlaps almost completely with the apical Rab7 compartment (Fig. 3G). We also analysed Snx16 function by generating cell clones with the loss of function allele *Snx16^Δ1^* (Rodal et al., 2011). In mutant cells, we detected DE-cadherin aggregates (arrowheads in Fig. 3E), which indicates a function of Snx16 in DE-cadherin trafficking. We confirmed that the DE-cadherin trafficking defect is indeed caused by the loss of *Snx16* by rescuing DE-cadherin accumulation with an Snx16 transgene (Fig. S2G). We conclude that Snx16 localises to Rab7 endosomes and is required for proper DE-cadherin trafficking.

Notably, *Snx16* mutant cells resemble *Rab7* mutants since DE-cadherin accumulation is modest and does not prevent ZA formation. A functional link between Snx16 and Rab7 is suggested by the finding that the DE-cadherin aggregates, which form in *Snx16* mutant cells, are detectable within the Rab7 compartment (arrowheads in Fig. 3F). This supports the idea that Snx16 is required to transport DE-cadherin out of the Rab7 compartment.

Since Rab7 is known to recruit retromer components to endosomes (Rojas et al., 2008; Seaman et al., 2009), we hypothesised that Rab7 also recruits Snx16 to endosomal DE-cadherin. If this is correct, increased amounts of Snx16 might suppress the need for Rab7 in recruiting Snx16, and thereby rescue DE-cadherin trafficking. To test this, we overexpressed Snx16 in *Rab7* mutant cell clones. This rescued DE-cadherin aggregation almost completely supporting the idea that Rab7 promotes DE-cadherin trafficking by recruiting Snx16 to endosomes (Fig. 3C,H).

We next asked whether the ability of Snx16 to rescue DE-cadherin aggregation is dependent on its capacity to form membrane tubules. In contrast to most sorting nexins, the tubulation activity of Snx16 is not dependent on a Bar domain but on a coiled-coil region (Wang et al., 2019). To test if Snx16 tubulation is required to rescue DE-cadherin trafficking we expressed Snx16*Δ*CC, an Snx16 protein in which the coiled-coiled domain had been deleted (Wang et al., 2019), in *Rab7* mutant cell clones. In contrast to wild type Snx16, Snx16*Δ*CC did not rescue DE-cadherin aggregation (Fig. 3D,H). This suggests that wild type Snx16 rescues DE-cadherin aggregation by forming membrane tubules.

The deletion in Snx16*Δ*CC does not affect its PX domain, which facilitates the interaction with DE-cadherin (Xu et al., 2017). This suggests that Snx16*Δ*CC is still able to associate with DE-cadherin. Consistently, we found that the DE-cadherin aggregates in *Rab7* mutant cells co-localise with Snx16*Δ*CC (arrowheads in Fig. 3D). We therefore propose that overexpressed Snx16*Δ*CC associates with DE-cadherin in *Rab7* clones, but is unable to facilitate the exit of DE-cadherin out of endosomes due to its inability to form tubules.

In summary, our data indicate that Snx16, which was previously shown to bind E-cadherin, localises to the Rab7 compartment. Loss of Snx16 leads to DE-cadherin aggregation within the Rab7 compartment. Loss of Rab7 also leads to DE-cadherin aggregation, but this defect can be rescued by increased levels of tubulation active Snx16. Together these data are consistent with a model, in which Rab7 recruits Snx16 to endosomes, where Snx16 binds DE-cadherin and forms tubules, which transport DE-cadherin out of the Rab7 compartment.

### Rab11 recruits Sec15 after DE-cadherin export from the Snx16 compartment

Our data suggest that Snx16 transports DE-cadherin out of the Rab7 compartment and indicate that the Rab7 and Rab11 compartments are connected. We therefore propose that Snx16 transports DE-cadherin from the Rab7 into the Rab11 compartment. This leads to the question of how DE-cadherin trafficking proceeds within the Rab11 compartment. One central function of Rab11 is the recruitment of its effector Sec15, a member of the exocyst complex, which is required for the fusion of vesicles with the PM (Prigent et al., 2003; Zhang et al., 2004). To localise where Rab11 recruits Sec15 we co-expressed HA-Rab11 and a mCherry tagged form of Sec15. The tag does not interfere with Sec15 function as revealed by the ability of Sec15-Cherry to rescue Sec15 depleted epithelia (Fig. S2I). Surprisingly, co-overexpression of Rab11 and Sec15 resulted in aggregates consisting of DE-cadherin, Rab11 and Sec15 (Fig. 4A). This indicates that a simultaneous increase of Rab11 and Sec15 levels interferes with DE-cadherin export out of the Rab11 compartment.

**Figure 4:**
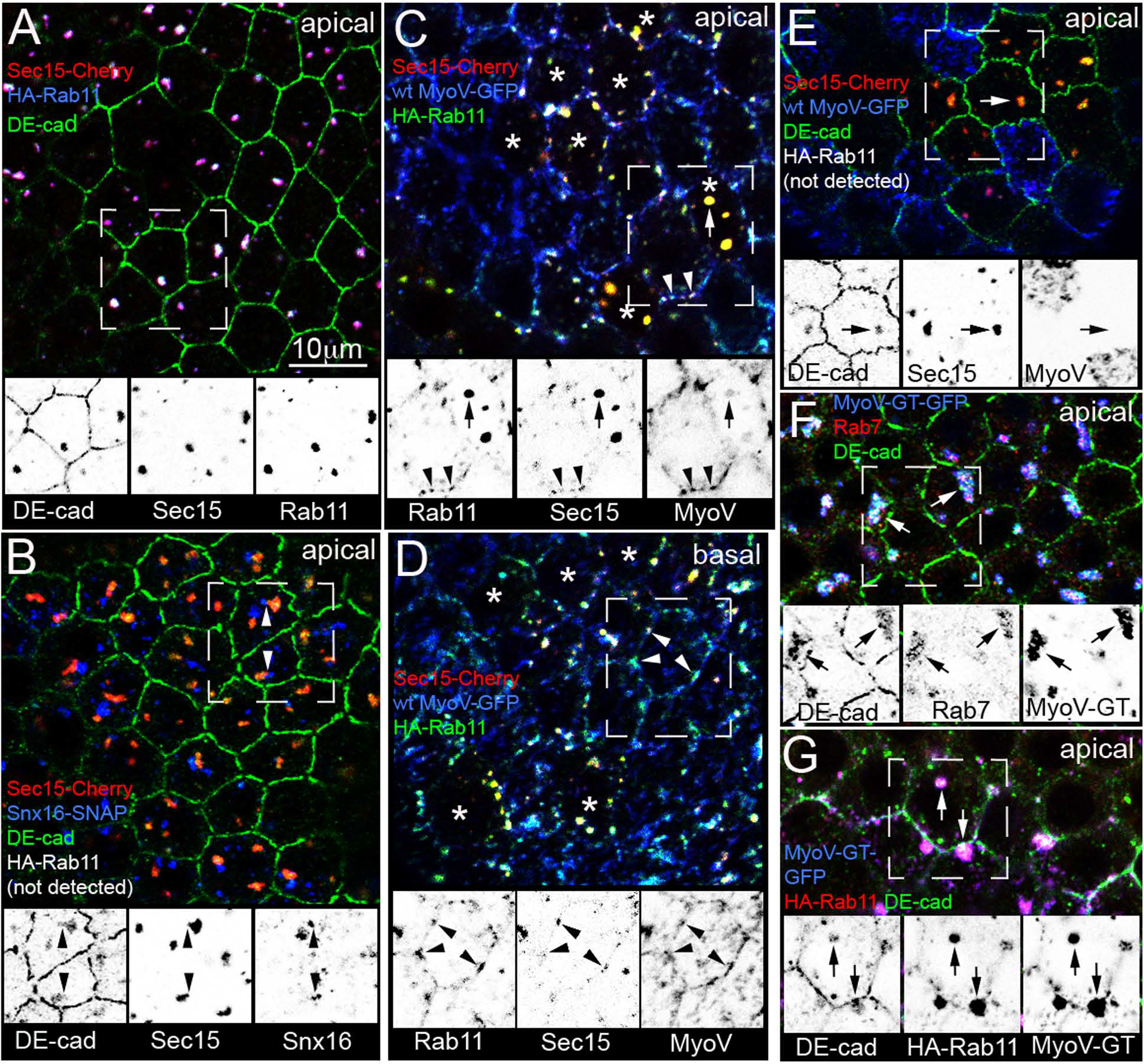
MyoV transports Rab11, Sec15 and DE-cadherin to the PM. Optical confocal sections through the follicular epithelium. Insets show single channels of the region marked by the dashed box. (**A**) Apical section of an HA-Rab11 (blue) and Sec15-Cherry (red) overexpressing epithelium that was stained for HA and DE-cadherin (green). HA-Rab11, Sec15-Cherry and DE-cadherin co-aggregate in the apical cytoplasm. (**B**) Apical section of an epithelium co-overexpressing wild type Snx16-SNAP (blue), HA-Rab11 and Sec15-Cherry (red) which was stained for SNAP and DE-cadherin (HA-Rab11 is not detected). DE-cadherin and Sec15 form aggregates. Snx16-SNAP localises next to these aggregates but does not overlap (arrowheads). (**C**) Apical section of an epithelium co-overexpressing HA-Rab11 (green), Sec15-Cherry (red) and MyoV-GFP (blue) stained for HA and GFP. Asterisks mark cells with low MyoV-GFP expression. Arrow indicates a Rab11/Sec15/MyoV aggregate in a cell with low MyoV expression. Rab11 and Sec15 signal intensities are strong, whereas MyoV is hardly detectable. Arrowheads indicate Rab11/Sec15/MyoV aggregates at the PM of a cell with a high MyoV-GFP expression level. At the PM the signal intensity of MyoV is stronger compared to the intensity in the arrow marked aggregate. By contrast, the intensities of the Rab11 and Sec15 signals at the PM are lower compared to the intensities in the arrow marked aggregate. Quantification of signal intensities of 20 cells from 5 different follicles: The signal intensity of Sec15 in the cytoplasmic aggregates is 10.88 times stronger than the signal intensity of Sec15 at the PM (±1.32). The signal intensity of Rab11 in the aggregates is 3.90 times stronger than the signal intensity of Rab11 at the PM (±0.60). The MyoV signal intensity at the PM is 13.89 times stronger than the signal intensity of MyoV in the aggregates (±3.58). (**D**) Basal section of the epithelium shown in (C). Arrowheads highlight co-localization of Rab11, Sec15 and MyoV in the basal region of the lateral membrane. (**E**) Apical section of an epithelium co-overexpressing HA-Rab11 (not detected), Sec15-Cherry (red) and MyoV-GFP (blue) that was stained for DE-cadherin (green). Sec15 and DE-cadherin co-aggregate in cells with low MyoV expression levels. Arrow points at a DE-cadherin/Sec15 aggregate in which MyoV is not detectable. (**F,G**) Epithelia expressing the dominant-negative form MyoV-GT-GFP (blue). The dominant-negative form of MyoV forms aggregates that co-localise with DE-cadherin (green). Rab7 (red, F) and HA-Rab11 (red, G) co-localise with the MyoV/DE-cadherin aggregates (100% for Rab7, n=21 follicles; 100% for HA-Rab11, n=5 follicles).

We mapped where DE-cadherin transport is blocked in relation to the apical endosomal Snx16 compartment. Notably, the blocked Rab11 compartment abuts the Snx16 compartment but hardly overlaps with it (arrowheads in Fig. 4B). This suggests a spatial separation of the Snx16 compartment from the compartment, where Rab11 accumulates with DE-cadherin and Sec15. We therefore propose that DE-cadherin associates with Rab11 and Sec15 only after DE-cadherin has left the Snx16 compartment. This raises the possibility that the binding of Snx16 to DE-cadherin prevents the recruitment of Rab11 (and Sec15) to DE-cadherin.

### MyoV transports DE-cadherin from the Rab11 compartment to the PM

A possible explanation for the block of DE-cadherin trafficking after simultaneous overexpression of Rab11 and Sec15 are insufficient levels of a factor, which transports the DE-cadherin/Rab11/Sec15 complex to the PM. A good candidate is the actin motor MyoV. Importantly, MyoV is another Rab11 effector, which also binds Sec15 (Jin et al., 2011). This raises the possibility that MyoV interacts with both, Rab11 and Sec15, during DE-cadherin transport. We examined if the three proteins co-localise by co-expressing a GFP-tagged form of MyoV with HA-Rab11 and Sec15-Cherry. We noted that the expression levels of MyoV-GFP varied significantly within the epithelium, which resulted in cells with high and very low MyoV-GFP expression. Such cell clone variations in Gal4/UAS induced gene expression had been reported previously (Skora and Spradling, 2010). MyoV-GFP co-localised in high and low areas with HA-Rab11 and Sec15-Cherry. However, the distribution pattern of the three proteins varied. Cells with low MyoV-GFP expression formed Rab11/Sec15 aggregates similar to cells without any MyoV-GFP expression (arrow in Fig. 4C). By contrast in cells with high MyoV-GFP expression, Rab11 and Sec15 aggregates were hardly detectable but when they did appear, they localised at or close to the PM (arrowheads in Fig. 4C). Notably, the highest MyoV-GFP intensity was detected at the PM. Rab11 and Sec15 were also detectable at the PM, albeit with lower signal intensities compared to the intracellular aggregates. A possible explanation for this protein distribution is that high MyoV-GFP levels rescue Rab11/Sec15 aggregation by delivering them to the PM. At the PM, Rab11 and Sec15 dissociate, whereas MyoV-GFP stays attached to the abundant F-actin at the PM. Low levels of MyoV-GFP are, on the other hand, not sufficient to rescue Rab11/Sec15 aggregation efficiently.

Consistent with a rescuing activity of MyoV-GFP, the strength of DE-cadherin aggregation was significantly reduced when MyoV-GFP was expressed in the Rab11/Sec15 overexpressing background. Cells with high MyoV-GFP levels formed almost no DE-cadherin aggregates, and cells with low MyoV-GFP formed DE-cadherin aggregates with low signal intensities (arrow in Fig. 4E). To quantify the MyoV-GFP rescuing activity we counted DE-cadherin aggregates in Rab11/Sec15 versus MyoV/Rab11/Sec15 expressing epithelia. To take into account that the signal intensity of the DE-cadherin aggregates in the MyoV/Rab11/Sec15 overexpressing epithelia is lower, we counted only those aggregates, whose signal intensity was higher than the DE-cadherin intensity measured at the PM. This revealed a tenfold reduction in the number of DE-cadherin aggregates in MyoV expressing cells, which declined from 3.1 (n=300 cells) to 0.3 (n=249 cells) aggregates per cell. Thus, MyoV-GFP suppresses the Rab11/Sec15 induced DE-cadherin aggregation. This indicates that DE-cadherin aggregates in Rab11/Sec15 overexpressing cells form because of insufficient MyoV levels.

We expressed a dominant-negative form of MyoV to examine if reduced MyoV activity affects DE-cadherin transport. This GFP-tagged C-terminal peptide encompasses the globular tail region (MyoV-GT-GFP), and perturbs the activity of the wild type protein by competing with it for cargos (Krauss et al., 2009; Wu et al., 1998). Consistent with previous reports, we observed big MyoV-GT-GFP aggregates in the follicular epithelium (Aguilar-Aragon et al., 2020). Notably, we also detected DE-cadherin within these aggregates (Fig. 4F,G). This indicates that dominant-negative MyoV interferes with DE-cadherin transport. HA-Rab11 and Rab7 also accumulated within the aggregates, which indicates that DE-cadherin trafficking is blocked within the endosomal system. Thus, dominant-negative MyoV interferes with endosomal sorting of DE-cadherin, which suggests that active MyoV is involved in transporting DE-cadherin out of endosomes.

### MyoV localises to an apical actin network and to basal actin bundles

To identify potential MyoV tracks for DE-cadherin transport we examined the distribution of the wild type form of MyoV-GFP in relation to the actin cytoskeleton. We detected MyoV puncta in an F-actin network in the apical area of the epithelium (arrow in Fig. 5A). This network is close to the endosomes, in which we detected endocytosed DE-cadherin (Fig. 1A). Consistently, also the apical Rab7 endosomes are in close proximity (Fig. 5C). The apical actin network is curved apically towards the centre of the cell (see Fig. S3A for an explanation of the geometry of the follicular epithelium and its cells). At the ZA, F-actin and DE-cadherin concentrate, which is consistent with the function of DE-cadherin in anchoring actin filaments (arrowheads in Fig. 5A). Notably, the MyoV-GFP signal also concentrates at the ZA. This together with MyoV-GFP puncta at the actin filaments and the fact that MyoV is an actin motor suggests a MyoV facilitated transport towards the ZA. Interestingly, the MyoV signal decreases in regions of the PM that are apical and basal to the ZA (green arrows and blue arrowheads in Fig. S3B-D). This supports the idea that the MyoV movement is specifically directed to the ZA. In summary, our data support a model, in which MyoV transports DE-cadherin vesicles from endosomes along the F-actin network to the ZA. This transport route is capable of stabilising the ZA by constantly delivering DE-cadherin from endosomes to the ZA.

**Figure 5:**
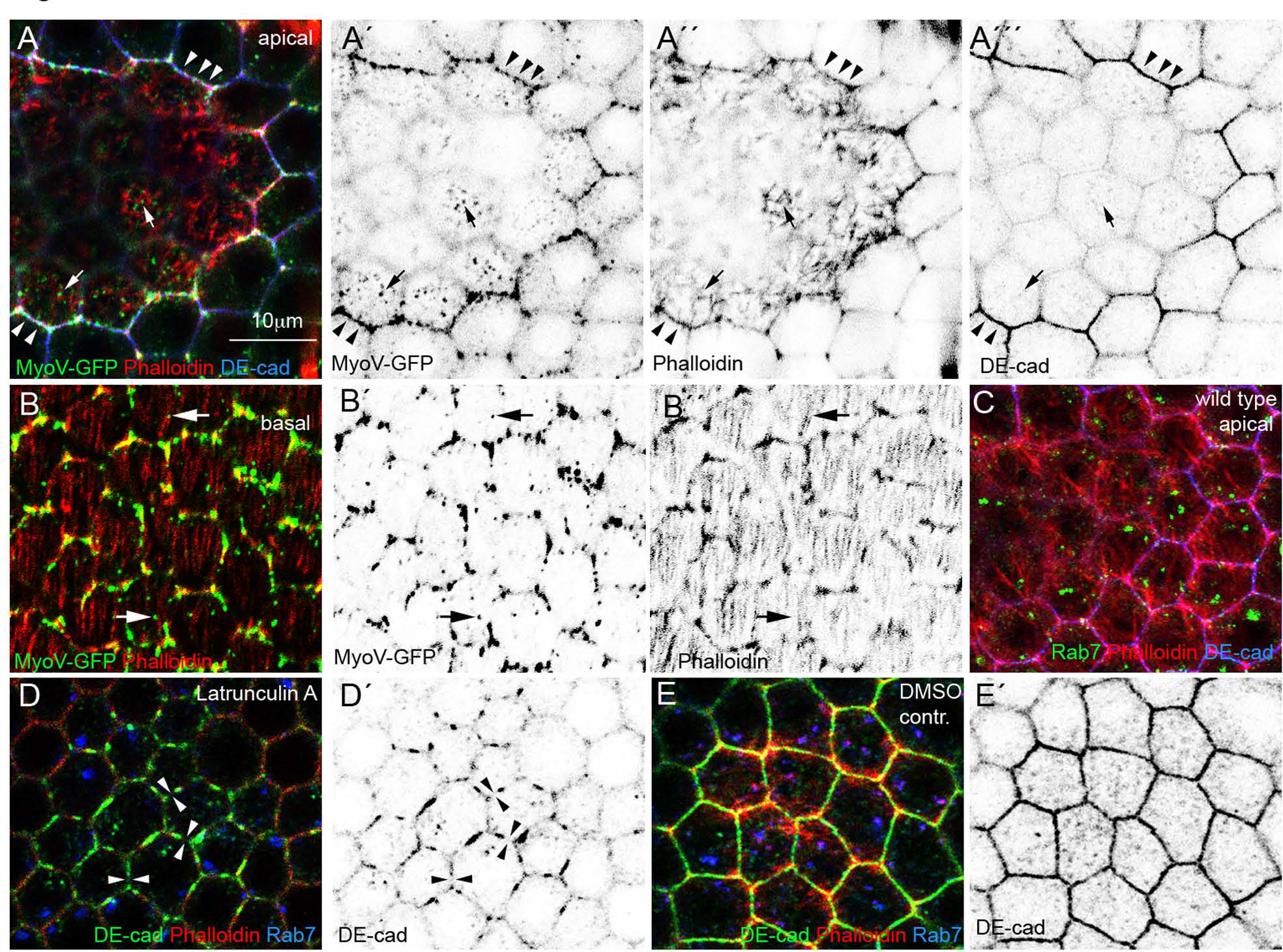
MyoV localises to apical and basal actin filaments. Apical (A) and basal (B) confocal sections of epithelia expressing MyoV-GFP (green) that are stained for Phalloidin (red). (**A**) The epithelium was also stained for DE-cadherin (blue). Single channels are shown in (Á-Á’’). Individual epithelial cells are cut at different positions along their apico-basal axis. This is due to the bending of the epithelium, which encapsulates an egg-shaped germline cyst. Cells in the middle show more apical views than the cells in the upper, lower and right parts of the picture. Within one cell, the apical actin network is curved from the ZA towards the centre of the cell (Fig. S3A shows a sketch explaining the geometry of the follicular epithelium and its cells). Arrows point at MyoV puncta that are close to actin filaments. Arrowheads mark the ZA, where the signal intensities of DE-cadherin, MyoV and Phalloidin are highest. (**B**) The basal section shows parallel arrays of actin filaments along which MyoV puncta are detectable (arrowheads). Single channels are shown in (B’-B’’) (**C**) Apical view of a wild type epithelium stained for Rab7 (green), Phalloidin (red) and DE-cadherin (blue). (**D**) Epithelium of a follicle that was treated for 2 hours with 20μM Latrunculin A and stained for DE-cadherin (green), Phalloidin (red) and Rab7 (blue). Arrowheads point at gaps within the ZA. Note that the F-actin is only detectable at the PM but not within the cell. (**D’**) shows DE-cadherin channel alone. (**E**) Epithelium of a control follicle for (D) that was treated in parallel with DMSO alone. F-actin is visible within the cytoplasm and the ZA is intact. (**É**) shows DE-cadherin channel alone. 82.38% (± 3.63) Latrunculin A treated cells have a fragmented PM (n=152 cells from 6 follicles), whereas only 2.55% (± 1.05) DMSO control cell show a fragmented ZA (n=287 cells from 9 follicles)

To test if the apical actin network is required for DE-cadherin transport, we treated ovaries with Latrunculin A, which causes rapid F-actin disassembly (Spector et al., 1983) and prevents actin polymerisation (Coue et al., 1987). As expected, Latrunculin A treatment disrupted the apical actin network completely. Notably, it also caused the fragmentation of the ZA (arrowheads in Fig. 5D). A possible explanation for this defect is that F-actin disruption interferes with DE-cadherin transport to the ZA, which leads to ZA fragmentation.

Interestingly MyoV-GFP, Sec15-Chery and HA-Rab11 are not only detectable apically but also in the basal region of the epithelium. Here, F-actin is organised in parallel bundles, which are reminiscent of stress fibres (Gutzeit, 1990). Like in the apical network, we detected MyoV-GFP puncta along the actin bundles (arrows in Fig. 5B). We also detected a high MyoV signal intensity in the basal region of the lateral PM, where MyoV co-localises with HA-Rab11 and Sec15-Cherry (arrowheads in Fig. 4D). This suggests that MyoV moves DE-cadherin/Rab11/Sec15 vesicles also along the basal actin bundles to the PM. Such a basal transport route for DE-cadherin vesicles represents an alternative exocytosis pathway, which could provide the pool of DE-cadherin that localises as puncta along the lateral membrane. In contrast to the ZA, DE-cadherin does not concentrate at the basal region but rather distributes along the lateral membrane. This suggests that DE-cadherin, which was secreted to the basal region is moved in an apical direction. This movement could be facilitated by a previously reported apically directed Cadherin flow, which has been observed in *Drosophila* (Woichansky et al., 2016) and in mammalian cells (Kametani and Takeichi, 2007).

### MyoV/DE-cadherin particles move towards the PM

To analyse MyoV facilitated transport of DE-cadherin in living follicles we used flies in which the coding region of mTomato was integrated into the DE-cadherin locus (Huang et al., 2009). Homozygous DE-cadherin:mTomato flies are viable indicating that the tagged DE-cadherin is functional. Consistently, DE-cadherin:mTomato localises like the wild type protein to the apical Rab7 compartment (arrows in Fig. 6A). Notably, DE-cadherin:mTomato also co-localises with MyoV-GFP in the apical actin network and along the basal stress fibres (arrows in Fig. 6B,C). This suggests that the tagged DE-cadherin forms transport vesicles with MyoV that move along F-actin.

**Figure 6:**
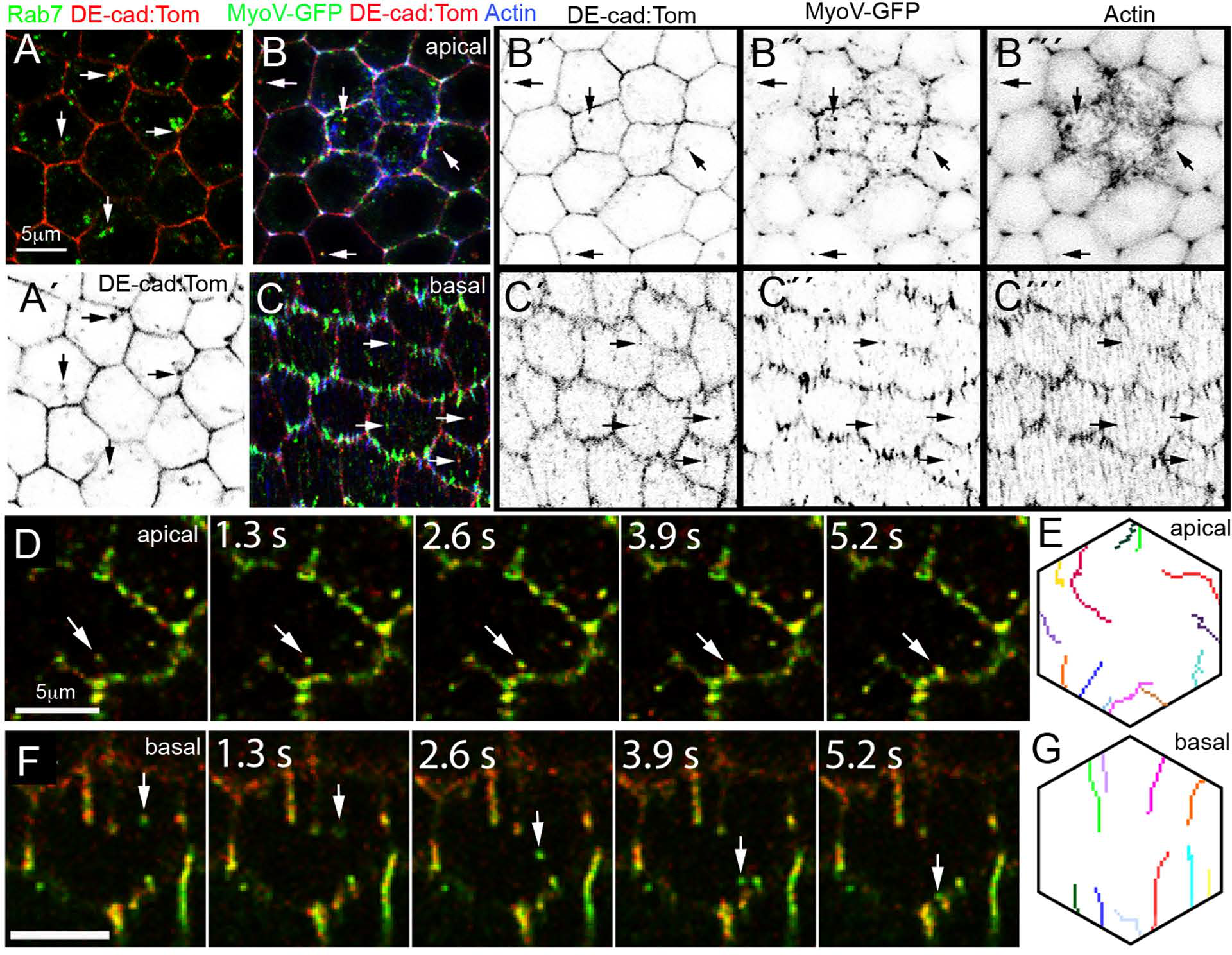
MyoV transports DE-cadherin particles towards the PM. (**A-C**) Optical confocal sections through the epithelium of fixed follicles. (**A**) Follicle expressing endogenously mTomato-tagged DE-cadherin (red) that was stained for Rab7 (green). Arrows point at DE- cadherin particles within the Rab7 compartment. 88.62% (± 3.47) of the Rab7 compartments co-localise with DE-cadherin particles (n=142 cells from 5 follicles). (**Á**) shows DE-cadherin channel alone. (**B,C**) Follicle expressing endogenously mTomato-tagged DE-cadherin (red) in which MyoV-GFP expression was induced and that was stained with Phalloidin (blue). Arrows point at DE-cadherin particles that co-localise with MyoV. The individual channels are shown to the right of the picture. (**B**) Apical section showing the area where the apical F- actin network is formed. (**C**) Basal section showing the parallel F-actin arrays. (**D,F**) Live imaging of follicles expressing endogenously mTomato-tagged DE-cadherin (red) in which MyoV-GFP (green) expression was induced. The arrows point at particles that are moving towards the PM. (**D**) Montage of movie S1 showing the follicular epithelium in an apical view. Five consecutive images of the movie are shown. (**F**) Montage of movie S2 showing a basal view. (**E,G**) Summaries of the trajectories of different apical (**E**) and basal (**G**) particles from different follicles projected onto one cell.

To test if the DE-cadherin:mTomato/MyoV-GFP puncta are indeed mobile we performed live imaging. We identified DE-cadherin:mTomato/MyoV-GFP particles in the apical and basal region of the epithelium that moved from central areas of the cell towards the PM (examples are shown in Fig. 6 D,F and in Movie 1,2). A summary of individual particle tracking paths revealed differences in the transport routes in apical and basal regions. Apical particles show more irregular paths (Fig. 6E), which is consistent with the network-like arrangement of the apical F-actin. By contrast, basal particles tend to move in a straight line towards the membrane (Fig. 6G). This reflects the basal arrangement of actin filaments in straight parallel bundles. These bundles align perpendicular to the anterior-posterior axis of the follicle (Gutzeit, 1990) (Fig. 6C’’ and Ć’’). Consistent with this organisation, basal particles did not move in all directions but only to opposite PMs, where the actin filaments presumably terminate. This is in contrast to the apical particles, which moved to all PMs. In summary, these data indicate that MyoV/DE-cadherin particles move along F-actin tracks towards the PM. This together with the findings that a dominant-negative MyoV results in cytoplasmic DE-cadherin aggregation and that disruption of F-actin causes ZA fragmentation strongly argues that MyoV transports DE-cadherin vesicles along F-actin to the PM.

In conclusion, our data suggest that MyoV transports DE-cadherin vesicles along the apical actin network to the ZA, and along the basal actin bundles to the basal-most region of the lateral PM. MyoV appears to use these two regions, where the epithelium provides F-actin tracks that run perpendicular to the apical-basal axis of the epithelium to secrete DE-cadherin to the lateral PM. Transport to the basal region of the lateral PM might act in combination with the Cadherin flow to supply the entire lateral PM with DE-cadherin, while the transport along the apical actin network provides DE-cadherin specifically for the ZA.

## Discussion

Our data suggest the following model for DE-cadherin trafficking (Fig. 7). DE-cadherin that was endocytosed from the lateral PM is transported to endosomes that localise to the apical region of the epithelium. Newly translated DE-cadherin is also transported to these apical endosomes, which leads to a confluence of the recycling and biosynthetic pathways. Our data suggest that Rab7 recruits Snx16 to these endosomes, which transports DE-cadherin via tubulation into the Rab11 domain. Rab11 orchestrates the formation of DE-cadherin transport vesicles by recruiting its effectors Sec15 and MyoV. The latter facilitates vesicle transport along an apical F-actin network towards the ZA. Sec15 recruits the other components of the exocyst complex, which are required for vesicle fusion with the PM (Grindstaff et al., 1998; Yeaman et al., 2004). This route reflects the previously proposed “apicolateral exocytosis” pathway (Woichansky et al., 2016), and stabilises the ZA by continuous DE-cadherin delivery.

**Fig. 7:**
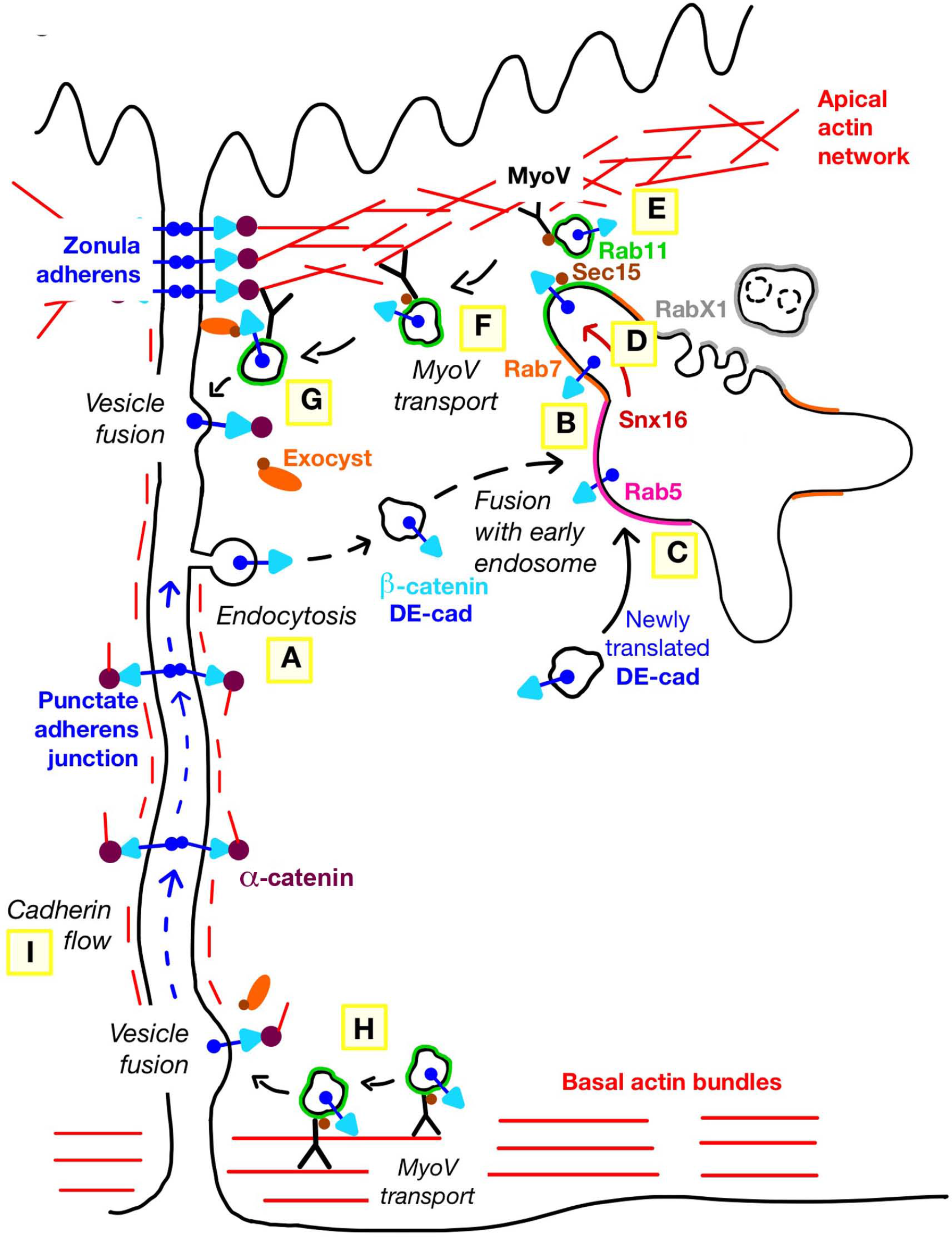
A model for DE-cadherin trafficking. Two epithelial cells are shown whose lateral PMs (left) are connected by belt-like adherens junctions forming the ZA and by punctate adherens junctions. An endosome is depicted within the right cell. The finger-shaped protrusions reflect different retrieval pathways. The degradative part where endolysosomes are formed is more vesicular (grey). Capitals indicate different steps in the DE-cadherin transport process to the PM: (**A**) DE-cadherin is endocytosed at the lateral PM. Vesicles with endocytosed (**B**) and newly translated (**C**) DE-cadherin fuse with the early endosome, which is limited by the Rab5 domain of the endosome (magenta). The endosome is subdivided into an endolysosomal part harbouring RabX1 (grey) and several branches into which proteins are sorted for recycling. (**D**) Within the endosome, Rab7 (orange) recruits Snx16, which moves DE-cadherin into the Rab11 domain (green). (**E**) Rab11 orchestrates the formation of DE-cadherin vesicles by recruiting its effectors Sec15 and MyoV. (**F**) MyoV moves DE-cadherin vesicles along an apical actin network to the ZA. (**G**) Sec15 and *β*-catenin cooperate in the recruitment of the exocyst complex, which is required for vesicle fusion with the PM. (**H**) DE-cadherin vesicles with Rab11, Sec15 and MyoV are also transported along basal F-actin arrays to the basal-most region of the lateral PM. (**I**) A cadherin flow distributes basally secreted DE-cadherin along the lateral membrane.

An additional pathway secretes DE-cadherin to the lateral PM, where the punctate adherens junctions form. Our data localise this pathway to the basal-most region of the epithelium, where parallel actin bundles provide tracks for the MyoV/DE-cadherin vesicles. We propose that after fusion of these vesicles with the PM, the apical membrane flow spreads DE-cadherin along the entire lateral PM to supply the punctate adherens junctions (Kametani and Takeichi, 2007; Woichansky et al., 2016). Our data show that Sec15 co-localises with Rab11 and MyoV in the basal region as well (Fig. 4D), which indicates that also the basal vesicles require the exocyst complex for their fusion with the PM. It remains to be determined whether these basal vesicles consist of newly synthesised or recycled DE-cadherin or both.

We revise our initial model for E-cadherin trafficking regarding the role of RabX1 (Woichansky et al., 2016). Initially, we proposed that RabX1 coordinates DE-cadherin recycling in endosomes. However, we recently found that RabX1 acts as an organiser of the degradative part of endosomes (Laiouar et al., 2020). Based on these findings and our current study we propose that DE-cadherin, which is targeted for degradation is sorted into the RabX1 domain of endosomes, where endolysosomal degradation starts. DE-cadherin destined for recycling is transported by Snx16 from the Rab7 into the Rab11 compartment.

Our data indicate that Rab7 and Snx16 are required for proper DE-cadherin sorting in endosomes, but mutant cells show only mild defects, which indicates that DE-cadherin secretion is not abolished. This is in contrast to Rab11 and exocyst mutants (Langevin et al., 2005; Woichansky et al., 2016). A possible explanation for the weaker defects is that DE-cadherin sorting is controlled by partially redundant mechanisms. The F-Bar proteins Cip4 and Nostrin (Zobel et al., 2015) and sorting nexins Snx1 (Bryant et al., 2007) and Snx4 (Solis et al., 2013) have been shown to be involved in E-cadherin sorting and are thus good candidates to support Rab7 and Snx16 in DE-cadherin transport to the Rab11 compartment.

A central protein for E-cadherin trafficking is *β*-catenin, which binds to the cytosolic part of E- cadherin. *β*-catenin is thought to mediate the association of the exocyst complex with E-cadherin vesicles by binding to the exocyst component Sec10 (Langevin et al., 2005). This *β*- catenin mediated DE-cadherin-exocyst linkage may be essential for the fusion of DE-cadherin vesicles with the PM. One reason for the loss of PM localisation of DE-cadherin in *β- catenin*/*armadillo* mutants could therefore be a failure in recruiting the exocyst complex to DE- cadherin vesicles A recent study reported that the apical localisation of Crumbs is also dependent on the exocyst complex and MyoV (Aguilar-Aragon et al., 2020). This suggests that DE-cadherin shares parts of its exocytosis pathway with other cargos. However, there must be significant differences in the sorting mechanisms of the two proteins. For instance, it seems unlikely that *β*-catenin links Crumbs to the exocyst complex, which raises the question of how the exocyst-Crumbs linkage is facilitated. Moreover, in contrast to DE-cadherin, Crumbs does not localise to the lateral membrane (Tepass et al., 1990) and is therefore unlikely to be transported along the basal stress fibres. Thus, mechanisms must exist preventing Crumbs vesicles from using the basal stress fibres. Also, the endosomal sorting mechanisms of DE-cadherin and Crumbs are different as Crumbs recycling was shown to be dependent on the retromer complex (Pocha et al., 2011).

Besides this exocyst facilitated secretion, other pathways must exist delivering integral membrane proteins to the PM. One example is the cell-cell adhesion protein Fasciclin2, whose localisation is undisturbed in *Rab11* mutant (Woichansky et al., 2016) and in exocyst depleted cells (Fig. S2J). It remains to be shown how Fasciclin2 exocytosis to the lateral membrane is regulated and whether newly translated Fasciclin2 is also sorted in endosomes.

Our study gives new insight into endosomal sorting of DE-cadherin, but many molecular mechanisms still have to be elucidated. One example is the activation of Rab11, which might be regulated by the TRAPII complex or by Parcas. Both factors have recently been shown to promote the GDP-GTP exchange that is necessary for Rab11 activation in *Drosophila* (Riedel et al., 2018). It is also unclear how exactly DE-cadherin/Sec15 vesicles are formed within the Rab11 compartment. The recently discovered FERARI complex is a dynamic machinery, which could orchestrate the recruitment of Rab11 to endosomes and the subsequent formation of DE-cadherin/Sec15/MyoV vesicles (Solinger et al., 2020). Moreover, it is unknown when DE-cadherin/Sec15 vesicles recruit the other components of the exocyst complex. The formation of the complex is a dynamic process involving the formation of subcomplexes (Ahmed et al., 2018). These subcomplexes could associate with E-cadherin either in the Rab11 compartment, when the vesicles move along the actin filaments or just before vesicle fusion with the lateral membrane. It will also be important to elucidate how exactly DE-cadherin vesicle fusion is controlled. A study in mammalian cells identified the polarity protein Par-3 as an exocyst receptor (Ahmed and Macara, 2017), which localises to the ZA in the *Drosophila* follicular epithelium (Franz and Riechmann, 2010). Par-3 is therefore an excellent candidate to initiate and maintain ZA formation by precisely targeting DE-cadherin vesicle fusion.

The conserved role of E-cadherin for epithelial homeostasis raises the question of to what extent our model for DE-cadherin trafficking can be applied to other organisms. The fact that RabX1 has been lost during vertebrate evolution (Klöpper et al., 2012) indicates changes in the degradative branch of endosomal sorting. By contrast, the interaction between Rab11, Sec15 and MyoV has been identified in yeast (Jin et al., 2011) suggesting that this very ancient trafficking module might have been adopted for E-cadherin transport when multicellular organisms have evolved. The fact that the human Rab11 orthologue Rab11a is required for PM localisation of E-cadherin (Desclozeaux et al., 2008; Lock and Stow, 2005) supports the idea that this transport module exists also in mammalian cells. The interaction between E- cadherin and Snx16 has also been demonstrated in human cells (Xu et al., 2017), which suggest that also endosomal retrieval mechanisms are conserved between flies and mammals. We therefore think that our model for DE-cadherin trafficking in the follicular epithelium provides a good framework to understand how changes in E-cadherin trafficking promote morphogenesis and how defective trafficking initiates disease processes.

## Materials and Methods

### Drosophila genetics

*Fly stocks*: *w; traffic jam*-Gal4 (provided by J. Brennecke), GR1-Gal4/TM3 Ser (provided by S. Roth), UASp-YFP-*Rab11* (Zhang et al., 2007)/TM3, w[*]; TM3, P(w[+mC]=UAS-shibire.K44A) 3-10/TM6B, Tb[1] (Bloomington Stock Centre), w; FRT40A *Rab5^2^*/Cyo (provided by A. Guichet), UAS-HA-*Rab11* (Woichansky et al., 2016) (insertions on second and third chromosome), w; y*Rab7* (Dunst et al., 2015) (provided by M. Brankatschk), w;; FRT82B *Rab11^dFRT^*/TM3Sb (Bogard et al., 2007) (provided by R.S. Cohen), FRT82B *Rab7^Gal4-knock- in^*/TM6 (Cherry et al., 2013) (provided by R. Hiesinger), UAS-wt*Snx16*-SNAP (Rodal et al., 2011), UAS-*Snx16ΔCC*-SNAP (Wang et al., 2019), UAS-GFP-*Snx16* (Wang et al., 2019), FRT42 *Snx16^Δ1^*/Cyo (Rodal et al., 2011) (all provided by A.A. Rodal), UAS-GFP-MyoV full length (Krauss et al., 2009) (provided by A. Ephrussi), ; UAS-GFP-MyoV-GT (Krauss et al., 2009) (provided by B.J. Thompson), UAS-*Sec15*-mCherry (Michel et al., 2011) (provided by C. Böckel), (y)w; DE-Cadherin:mTomato (Huang et al., 2009) (provided by S. Bogdan), *Sec15* RNAi (VDRC, KK 105126) .

Flies were raised at 27°C and dissected 1-3 days after hatching. The only exception is the experiment with dominant-negative *shibire* flies, which were raised at 18°C to avoid lethality. After hatching flies were kept at 27°C for 24 hours to induce expression and then dissected.

All UAS-transgenes were induced with *traffic jam*-Gal4 with the exceptions of the experiments shown in Fig. 1D and Fig. S2G where GR1-Gal4 was used to induce UAS-YFP*- Rab11* and UAS-GFP*-Snx16*.

*Recombination of Rab11^dFRT^ allele with FRT82: Rab11^dFRT^* was initially on a chromosome harbouring the FRT site next to the mutation in *Rab11* (Bogard et al., 2007). To use the conventional *FRT82B* for clone induction we recombined the *Rab11^dFRT^* allele with the FRT82B. This removed a cell cycle affecting mutation in the background.

*Detailed experimental genotype:*

Figure 1B: w; *traffic jam*-Gal4 Sp/+; UAS-HA*-Rab11*/+; Figure 1C, 1F and 1G: w; *traffic jam*- Gal4 Sp/+; UAS-YFP*-Rab11*/+; Figure 1D: hsFLP/w; FRT40A *Rab^2^*/FRT40A RFP; UAS-YFP-*Rab11*/GR1-Gal4; Figure 1H: w; *traffic jam*-Gal4 Sp/+; UAS-YFP- *Rab11*/UAS-HA-*Rab11;* Figure 1I: w; *traffic jam*-Gal4 Sp/+; UAS-YFP-*Rab11*/UAS- *DNshibire;* Figure 2A: w; *traffic jam*-Gal4 Sp/+; UAS-YFP-*Rab11*/+; Figure 2D: w; *traffic jam*-Gal4 Sp/+; UAS-HA*-Rab11*/y*Rab7;* Figure 2F, 2G and 2H: hsFLP/w;; FRT82B *Rab11*/FRT82B RFP; Figure 3A and 3B: hsFLP/w;; FRT82B *Rab7*/FRT82B RFP; Figure 3C: hsFLP/w; *traffic jam*- Gal4 UAS-*Snx16wt*-SNAP/+; FRT82B *Rab7*/FRT82B RFP; Figure 3D: hsFLP/w; *traffic jam*-Gal4 UAS-*Snx16ΔCC*-SNAP/+; FRT82B *Rab7/*FRT82B RFP; Figure 3E and 3F: hsFLP/w; FRT42 *Snx16^Δ1^*/FRT42 RFP; Figure 3G: w; *traffic jam-*Gal4 UAS-*Snx16wt*- SNAP/Cyo; Sb/TM3Ser Figure 4A: w; *traffic jam-*Gal4 Sp/+; UAS-HA*-Rab11/*UAS-*Sec15*-mCherry; Figure 4B: w; *traffic jam*-Gal4 UAS-*Snx16wt-*SNAP/+; UAS-HA*-Rab11/*UAS-*Sec15*-mCherry; Figure 4C-E: w; *traffic jam*-Gal4 Sp/+; UAS-*Sec15*-mCherry UAS-*MyoV-FL*-GFP/UAS*-*HA-*Rab11;* Figure 4F: w; *traffic jam*-Gal4 Sp/+; UAS-*MyoV-GT-*GFP/+; Figure 4G: w; *traffic jam-* Gal4 Sp/+; UAS-*MyoV-GT*-GFP/ UAS-HA-*Rab11*; Figure 5A and 5B: w; *traffic jam-*Gal4 Sp/+; UAS-*MyoV-FL*-GFP/+ Figure 6A: w; *traffic jam*-Gal4 *DE-Cad*:Tom/Cyo; Sb/TM3Ser; Figure 6B, 6C, 6D and 6F: w; *traffic jam*-Gal4 *DE-Cad:*Tom/Cyo; UAS-*MyoV-FL*-GFP/+; Figure S1: *w; traffic jam-*Gal4 Sp/+; UAS-YFP-*Rab11*/+ Figure S2A: hsFLP/w;; FRT82B *Rab11*/FRT82B RFP; Figure S2B: hsFLP/w; *traffic jam-*Gal4 Sp/UAS-HA*-Rab11;* FRT82B *Rab11*/FRT82B RFP; Figure S2C and S2D: *w; traffic jam-* Gal4 Sp/+; UAS-HA*-Rab11*/y*Rab7;* Figure S2F: hsFLP/w; FRT42 *Snx16^Δ1^*/FRT42 RFP; Figure S2G: hsFLP/w; FRT42 *Snx16^Δ1^*/FRT42 RFP; UAS-GFP-*Snx16*/GR1-Gal4; Figure S2H and S2J: w; *traffic jam*-Gal4 Sp/+; *Sec15* RNAi/+; Figure S2I: w; *traffic jam-* Gal4 Sp/+; *Sec15* RNAi/UAS-*Sec15*-mCherry; Figure S3B, S3C and S3D: w; *traffic jam-*Gal4 Sp/+; UAS*-MyoV-FL*-GFP

### RNAi induction and the generation of genetic mosaics

For RNAi knockdown in the follicular epithelium, UAS-inducible RNAi transgenes were driven with *traffic jam*-Gal4.

Genetic mosaics were generated by using a heat-shock (hs) inducible Flipase (FLP). To induce the hs-FLP for generation of homozygous mutant cell clones heat shocks were applied during pupal stages by placing the vials in a 37 °C water bath for 1 hour once a day until hatching. Flies were raised at 27°C and females were dissected 24 to 48 h after hatching.

### Immunohistology

Ovaries were dissected in Schneider’s medium and fixed in 4% formaldehyde in PBS for 10 minutes at room temperature. Ovaries were washed and permeabilized with 0.1% TritonX- 100 in PBS. Next, ovaries were blocked with 0.5% BSA in 0.1% TritonX/PBS containing 0.5

µg/ml DAPI for nucleus staining for 20 minutes. Both primary and secondary antibodies were diluted in 0.1% Triton/PBS and incubated for 3 hours at room temperature. Ovaries were washed and mounted in Vectashield.

Primary antibodies were used at the following concentrations: rat anti-DE-cadherin 1:50 (DSHB, DCAD2), mouse anti-Armadillo 1:100 (DSHB, N2 7A1), goat anti-Golgin245 (Riedel et al., 2016) 1:500 (DSHB), rabbit anti-GM130 1:100 (Abcam, ab30637), mouse anti-Rab7 (Riedel et al., 2016) 1:20 (DSHB), rabbit anti-Rab5 1:500 (Abcam, ab31261), goat anti-GFP FITC 1:100 (GeneTex, GTX26662-100), mouse anti-HA 1:100 (Santa Cruz Biotechnology, 11867423001), rabbit anti-RFP 1:200 (Rockland, 600-401-379), rabbit anti-aPKC 1:200 (Santa Cruz Biotechnology, sc-216)

Images were acquired using Leica TSC SP5 confocal microscope equipped with HyD detectors using 63x/HCX PL APO 1.3 glycerol immersion objective (Leica) at a resolution of 1,024 × 1,024. If necessary, images adjusted for gamma. Images were edited and assembled with Adobe Photoshop CS3.

### Pulse-chase endoassay

Living ovaries were incubated at room temperature in a pulse-chase solution containing anti-DE-cadherin antibody (1:25), 10% FCS and 0,2 mg/ml Insulin in Schneider’s medium. Incubation was performed for 2 minutes on a rocking platform. Then the ovaries were washed and incubated in a solution containing only 10% FCS and 0,2 mg/ml Insulin in Schneider’s medium for different time periods. Ovaries were washed and fixed in 4% formaldehyde. Staining was performed as described.

### SNAP staining

Living ovaries were incubated at room temperature in SNAP solution containing 3 μM SNAP-Cell TMR-Star or SNAP-Cell 647SiR (New England BioLabs, S9105S and S9102S), 10% FCS and 0,2 mg/ml Insulin in Schneider’s medium. Incubation was performed for 30 minutes on a rocking platform. Ovaries were washed and fixed in 4% formaldehyde. Staining was carried on as described. All steps were performed protected from light.

### Lysotracker staining

Living ovaries were incubated at room temperature in Lysotracker solution containing 10 μM LysoTracker^TM^ Red (Thermo Fisher Scientific, L7528), 10% FCS and 0,2 mg/ml Insulin in Schneider’s medium. Incubation was performed for 30 minutes on a rocking platform. Ovaries were washed and fixed in 4% formaldehyde. Staining was carried on as described. All steps were performed protected from light.

### Phalloidin staining

Ovaries were dissected in Schneider’s medium and fixed in 4% formaldehyde in PBS for 10 minutes at room temperature. Ovaries were washed and permeabilized with 0.1% TritonX-100 in PBS. Next, ovaries were blocked with 0.5% BSA in 0.1% TritonX/PBS containing 0.5 µg/ml DAPI for nucleus staining for 20 minutes. Additionally, Alexa Fluor^TM^ 568 Phalloidin (Thermo Fisher Scientific, A12380) was added to the blocking solution in a dilution of 1:100 (400x stock solution). Both primary and secondary antibodies were diluted in 0.1% Triton/PBS and incubated for 3 hours at room temperature. After incubation with both primary and secondary antibodies, ovaries were washed and mounted in Vectashield.

### Latrunculin A treatment

Ovaries were dissected in Schneideŕs medium and then treated with 20 μM Latrunculin A (Sigma-Aldrich, L5163) in Schneideŕs medium containing 10% FCS and 0.2 mg/ml Insulin for 2 hours at the room temperature on a rocking platform. Ovaries were washed and fixed in 4% formaldehyde. Staining was performed as described.

### Image analysis and quantification

All pictures shown in the Figures are single confocal sections. We usually stain five to eight ovaries in one experiment, and each experiment was performed at least twice. We took images according to the following procedure: 7-9 images are taken along the apical-basal axis of the epithelium at a distance of 0.8 μm, which were subsequently analysed for possible defects. We took the ZA as a landmark, which we recognised by the high intensity of the DE-cadherin staining at the PM. An additional sagittal section was taken to determine the developmental stage of the follicle.

Image quantification was performed before “gamma adjustment”. Some quantifications (indicated below) were done with maximal projections of z-stacks.

Quantification of the overlap between different cell compartments and/or proteins (Figures 1A, 1B, 1C, , 1D, 1I, 2D, 2E, 2F-I, 3B, 3D, 3F, 3G, 4F, 4G, 6A and S2C-E). Using the “Polygonal

Lasso” tool of Photoshop CS3 an area showing the ZA was selected and cropped for further analysis. In ImageJ/FIJI with Plugins>Analyse>Cell counter tool, the overlap and the total number of the compartments was counted. The ratio of the compartments/proteins overlapping and the total number of the compartment was calculated. The analysis was performed with the confocal single sections, except for the Figures 1D, 2F-I, 3B, 3D and 3F, where the snapshots of maximal projections of the z-stacks were analysed. Data for Figures 2I and S2E were analysed and presented using Graph Pad Prism Software.

*Quantification of the size of Rab7 compartments (Figures 2B and 2C).* A specific area within a follicle was selected and cropped for further analysis. In ImageJ/FIJI with the “Freehand selection” tool, the size of Rab7 compartments were selected and the size was measured using the Analyse>Measure tool. The minimal threshold size for the Rab7 compartment was set to 0.48 µm^2^. The values were saved in ROI manager (Analyse>Tools>ROI manager) and exported to the Excel file, where the average size of Rab7 compartments was calculated. The analysis was performed with the confocal single sections.

Quantification of the number of DE-cadherin aggregates in Rab7, Snx16 and Rab11 clones and in Rab7 clones overexpressing Snx16 forms (Figures 3A, 3C, 3D, 3E, 3H, S2B and S2G): The ratio between the number of cell clones showing the DE-cadherin aggregation and the total cell clones counted was calculated. Data for Fig. 3H were analysed and represented using Graph Pad Prism Software. The analysis was performed with the snapshots of maximal projections of the z-stacks.

Quantification of signal intensities at the plasma membrane and in the cytoplasm in MyoV-GFP/Sec15-Cherry/HA-Rab11 expressing ovaries (Figure 4C). Using the “Polygonal Lasso” tool, an area with two cells was selected, one with low MyoV expression and one with high MyoV expression. In ImageJ/Fiji with the “Straight line” tool, a line was drawn along the membrane (in a cell with high MyoV expression) and along the aggregate in the cytoplasm (in a cell with low MyoV expression). The signal intensity values for MyoV, Sec15 and Rab11 compartments, both along the membrane and in the cytoplasm were obtained by Analyse>Plot Profile. The values were exported to the Excel sheet and the average signal intensity values for each membrane and aggregate in the cytoplasm were calculated. Finally, the ratio between the intensity values in the cytoplasm/aggregate and on the membrane for each compartment was calculated. The analysis was performed with the confocal single sections.

Quantification of the intensity of DE-cadherin aggregates with regard to DE-cadherin signal intensity at the PM in HA-Rab11/Sec15-Cherry versus MyoV-GFP/HA-Rab11/Sec15-Cherry expressing epithelia (Figure 4E): An area containing a minimum of 20 cells was selected and cropped for further analysis. In ImageJ/FIJI, the “Freehand line” tool was selected, the plot intensity graph for the two faintest membranes was obtained using Analyse>Plot Profile tool. The values were exported to the Excel file and the intensity average was calculated. The calculated average intensity was used as a reference value and set as the maximum in Image>Adjust>B&C. The DE-cadherin aggregates were counted using Plugins>Analyse>Cell counter tool. Finally, the exact number of cells was counted and the number of DE-cadherin aggregates per cell was calculated. The analysis was performed with the snapshots of maximal projections of the z-stacks.

Quantification of the number of fragmented membranes in Latrunculin A and DMSO treated follicles (Figure 5D and 5E). An area showing the ZA was selected and cropped for further analysis. In ImageJ/FIJI, Plugins>Analyse>Cell counter tool was used to count the cells that had a fragmented ZA. Finally, the percentage of cells having fragmented a ZA after treatment with Latrunculin A and DMSO was calculated. The analysis was performed with the confocal single sections.

### Live Imaging

Female flies were kept at 27°C for 1-3 days on yeasted fly food after hatching. Ovaries were dissected in Schneideŕs medium and single ovarioles extracted using Dumont forceps no. 5 (FineScience) as described (Prasad et al., 2007). Ovarioles were transferred to X-well chambers (Sarstedt 94.6190.802) equipped with 200 µl live imaging-medium (Schneideŕs *Drosophila* medium containing 15% FCS and 0.2 mg/ml insulin). Egg chambers were imaged on an inverted Leica LSM SP5 confocal microscope equipped with HyD detectors using a 63x/HCX PL APO 1.3 glycerol immersion objective (Leica). Time-lapse videos were obtained in 1.3 s time intervals at a resolution of 512 × 256.

For illustration of individual apical and basal tracks, the final track of an image sequence was selected and re-colored using the Wand (Tracing) tool, Color Picker and Flood Fill tool. Tracks were saved individually and then assembled in a schematic epithelial cell in Adobe Illustrator 2022 according to size.

## Acknowledgements

We thank the Bloomington and VDRC stock centres, Teresa Bonello, Anne Ephrussi, Sonia Lopez de Quinto, Barry Thompson, Robin Hiesinger, Avital Rodal and Christian Bökel for fly stocks, the Developmental Studies Hybridoma Bank for antibodies, Nadine Kraft for technical assistance and Carlos Riechmann for discussions, Georg Stoecklin and Martin Schmelz for sharing their lab space with us. We acknowledge the support of the Core Facility Platform Mannheim and in particular CF LIMa, Live Cell Imaging Mannheim (DFG INST 91027/9-1 FUGG). This work was funded by grants from the German Cancer Aid (Deutsche Krebshilfe) and the German Research Foundation (DFG, RI 1225/2-2).

## Authors contributions

D.T. and N.B. performed the experiments and V.R. conceived the project, raised funding and wrote the manuscript.

## Supplementary Figures

**Fig. S1:**
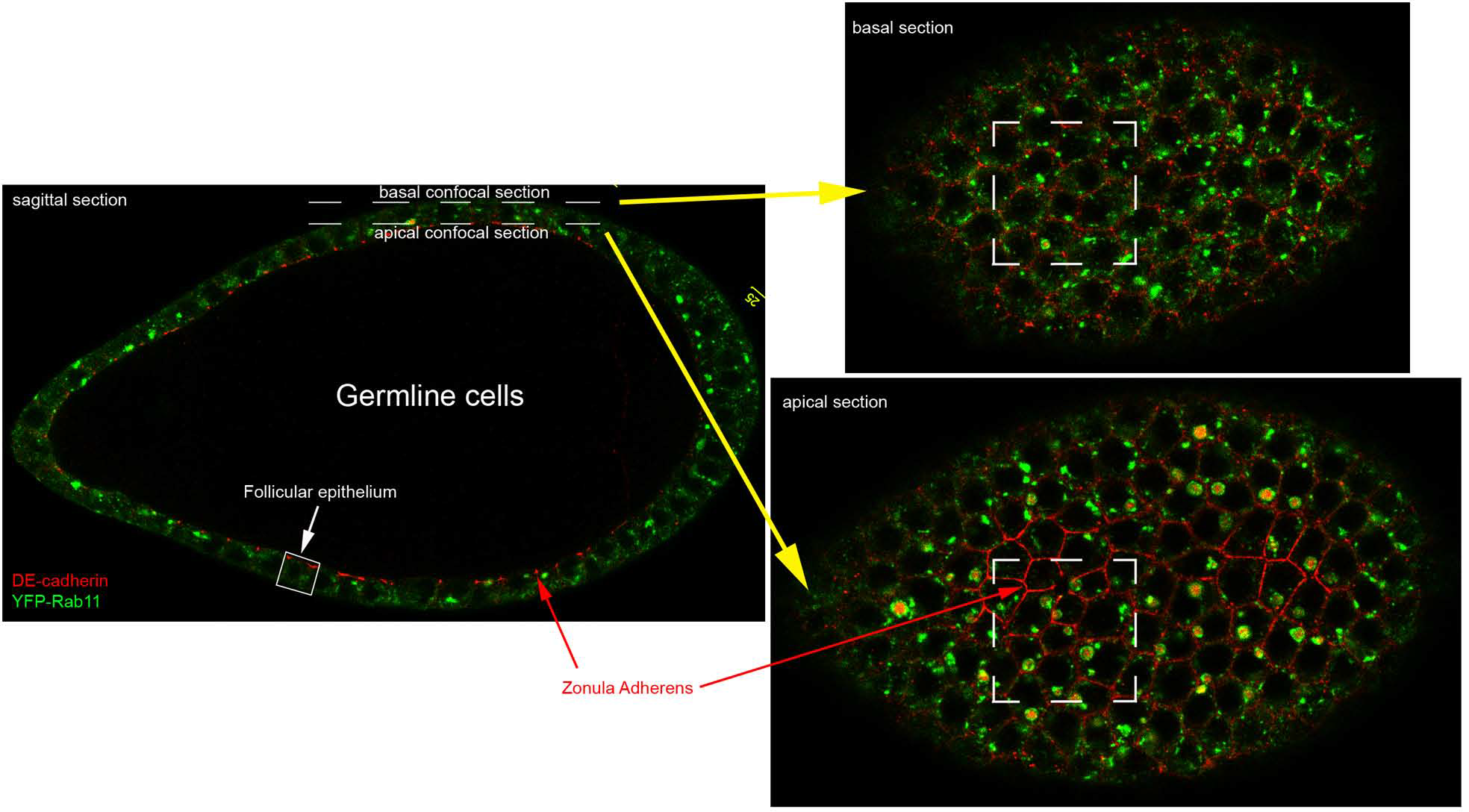
Analysis of protein distribution in the follicular epithelium using confocal pictures. The left picture shows a sagittal confocal section of a follicle stained for DE-cadherin (red) and YFP-Rab11 (green). Right pictures show a basal (up) and an apical (down) confocal section that was taken in regions corresponding to the dashed white line in the left picture.

**Fig. S2:**
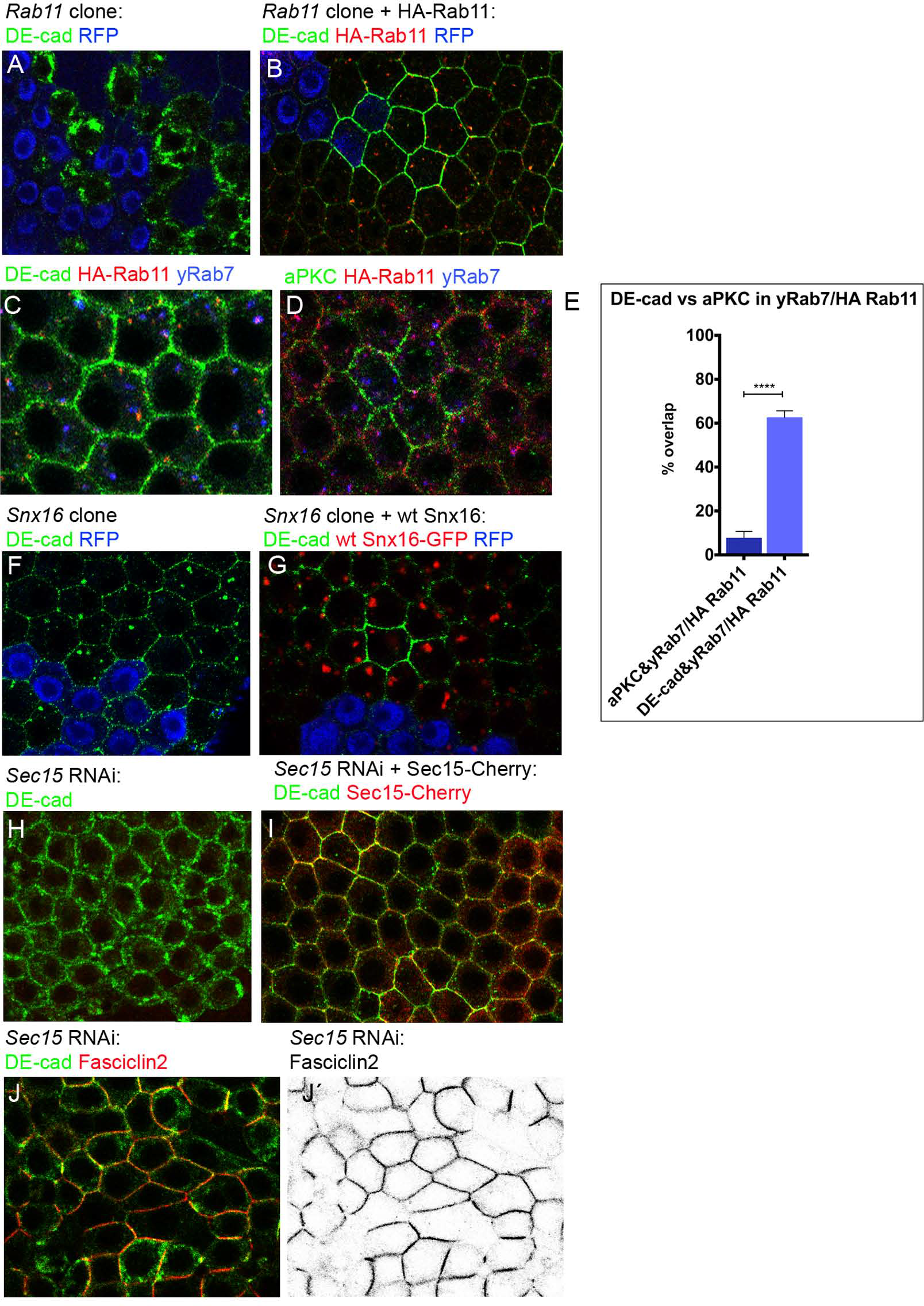
Control experiments. (**A**) *Rab11* mutant cell clone marked by the absence of RFP (blue) stained for DE-cadherin (green). Strong DE-cadherin accumulation is detectable (**B**) *Rab11* mutant cell clone in an epithelium in which HA-Rab11 (red) was expressed. DE- cadherin (green) localises to the PM and intracellular DE-cadherin accumulation is completely rescued by HA-Rab11 (red) indicating wild type activity of HA-Rab11 (100%, n=6 clones). (**C**) HA-Rab11 expressing follicle (red) stained for DE-cadherin (green) and Rab7 (blue). (**D**) Control experiment with an HA-Rab11 expressing follicle (red) stained for DE-cadherin (green) and aPKC (blue). (**E**) Quantification of the overlaps of the yRab7/HA-Rab11 compartment with DE-cadherin and aPKC. Data are shown as mean ± SEM. Two-tailed t-test (equal variance, *α* = 0.05) was performed and p values are presented as ****p < 0.0001 (compared groups: aPKC and yRab7/HA Rab11 overlap versus DE-cad and yRab7/HA Rab11 overlap). The quantification was performed with 5 follicles (n=199 cells) for the aPKC and yRab7/HA Rab11 and with 6 follicles (n=324 cells) for DE-cad and yRab7/HA Rab11. (**F**) *Snx16* mutant cell clone marked by the absence of RFP (blue) stained for DE-cadherin (green). *Snx16* mutant cells form small DE-cadherin aggregates. (**G**) *Snx16* mutant cell clone in an epithelium in which Snx16-GFP (red) was expressed. DE-cadherin (green) aggregation is rescued (92%, n=12 clones). (**H**) Epithelium in which Sec15 was depleted by expression of a UAS-Sec15 RNAi construct and that was stained for DE-cadherin (green). Note the strong intracellular accumulation of DE-cadherin. (**I**) Co-expression of UAS-Sec15 RNAi and a UAS-Sec15- Cherry constructs. Expression of UAS-Sec15-Cherry (red) rescues DE-cadherin (green) intracellular DE-cadherin aggregation and promotes PM localisation of DE-cadherin (92% n=13 follicles). The analysis of the rescue was performed with the single confocal sections. (**J**) Epithelium in which Sec15 was depleted by RNAi and that was stained for DE-cadherin (green) and Fasciclin2 (red). DE-cadherin accumulates within the cell, whereas Fascilin2 still to the PM is not affected. (**J’**) Fasciclin2 channel alone.

**Fig. S3:**
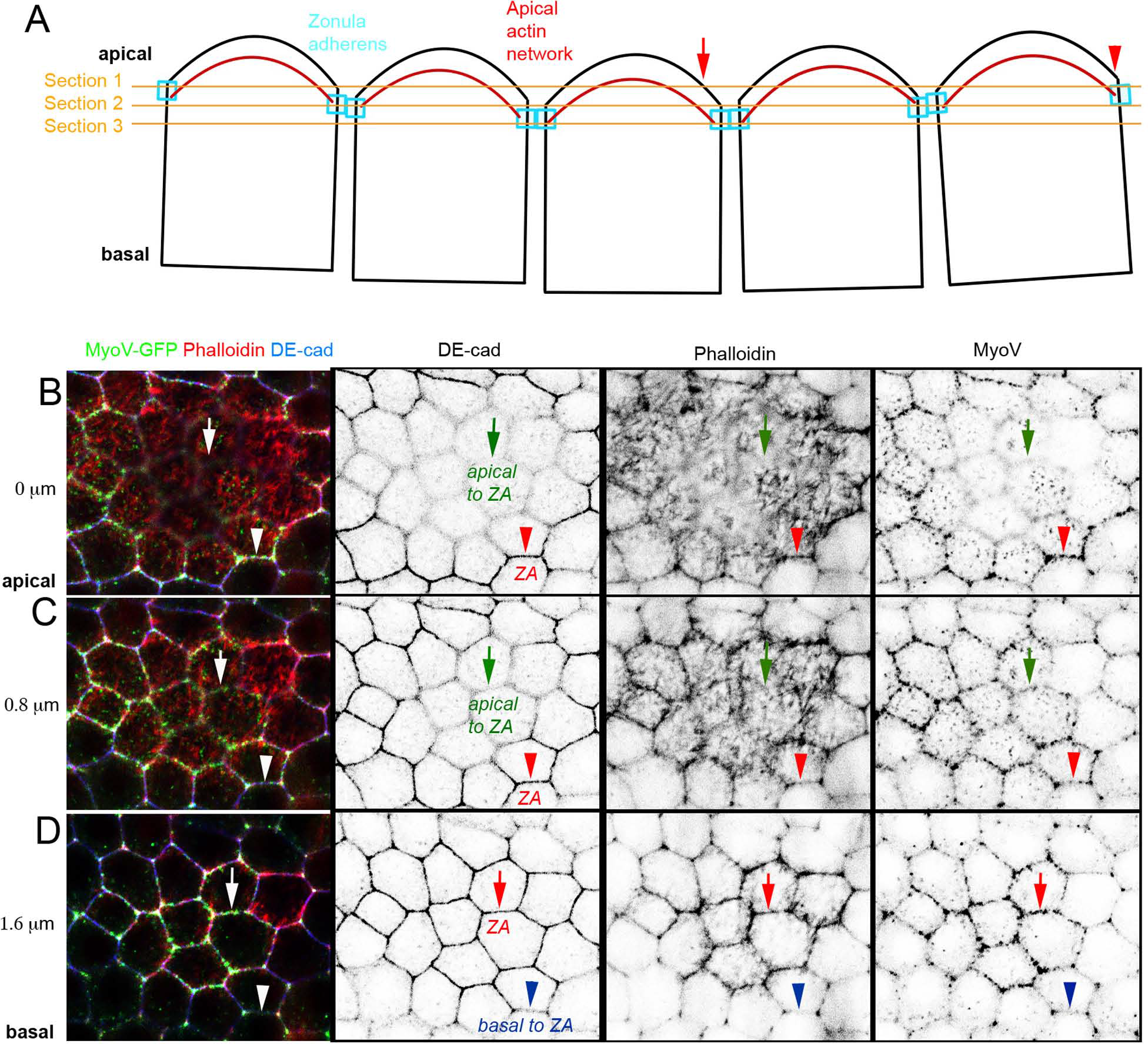
MyoV concentrates at the ZA. (**A**) Sketch depicting the geometry of the follicular epithelium and its cells. The epithelial monolayer is slightly bent due to the egg-shaped germline cyst. The apical actin network (red) is anchored at the ZA (blue) and curved towards the centre of the cell. The level of the three optical sections shown in (B-D) is indicated by the orange lines. (**B-D**) Images showing the apical region of the epithelium in three different planes. The spacing of the planes is 0.8 μm. The epithelium is stained for MyoV-GFP (green), Phalloidin (red) and DE-cadherin (blue). Arrows and arrowheads follow the borders of two cells from apical to basal showing how the distribution of DE-cadherin, F-actin and MyoV changes.

## Legend for Movies

**Movie 1**: The movie shows cells expressing endogenously mTomato-tagged DE-cadherin (red) and *traffic jam*-Gal4 induced MyoV-GFP (green) in the apical area of the epithelium.

**Movie 2**: The movie shows cells expressing endogenously mTomato-tagged DE-cadherin (red) and *traffic jam*-Gal4 induced MyoV-GFP (green) in the basal area of the epithelium.

## References

1. Aguilar-Aragon, M., G. Fletcher, and B.J. Thompson. 2020. The cytoskeletal motor proteins Dynein and MyoV direct apical transport of Crumbs. Dev Biol. 459:126–137.

2. Ahmed, S.M., and I.G. Macara. 2017. The Par3 polarity protein is an exocyst receptor essential for mammary cell survival. Nat Commun. 8:14867.

3. Ahmed, S.M., H. Nishida-Fukuda, Y. Li, W.H. McDonald, C.C. Gradinaru, and I.G. Macara. 2018. Exocyst dynamics during vesicle tethering and fusion. Nat Commun. 9:5140.

4. Bogard, N., L. Lan, J. Xu, and R.S. Cohen. 2007. Rab11 maintains connections between germline stem cells and niche cells in the *Drosophila* ovary. Development. 134:3413–3418.

5. Brüser, L., and S. Bogdan. 2017. Adherens Junctions on the Move-Membrane Trafficking of E-Cadherin. Cold Spring Harb Perspect Biol. 9.

6. Bryant, D.M., M.C. Kerr, L.A. Hammond, S.R. Joseph, K.E. Mostov, R.D. Teasdale, and J.L. Stow. 2007. EGF induces macropinocytosis and SNX1-modulated recycling of E- cadherin. J Cell Sci. 120:1818–1828.

7. Cadwell, C.M., W. Su, and A.P. Kowalczyk. 2016. Cadherin tales: Regulation of cadherin function by endocytic membrane trafficking. Traffic. 17:1262–1271.

8. Cherry, S., E.J. Jin, M.N. Ozel, Z. Lu, E. Agi, D. Wang, W.H. Jung, D. Epstein, I.A. Meinertzhagen, C.C. Chan, and P.R. Hiesinger. 2013. Charcot-Marie-Tooth 2B mutations in rab7 cause dosage-dependent neurodegeneration due to partial loss of function. Elife. 2:e01064.

9. Choi, J.H., W.P. Hong, M.J. Kim, J.H. Kim, S.H. Ryu, and P.G. Suh. 2004. Sorting nexin 16 regulates EGF receptor trafficking by phosphatidylinositol-3-phosphate interaction with the Phox domain. J Cell Sci. 117:4209–4218.

10. Classen, A.K., K.I. Anderson, E. Marois, and S. Eaton. 2005. Hexagonal packing of *Drosophila* wing epithelial cells by the planar cell polarity pathway. Dev Cell. 9:805–817.

11. Coue, M., S.L. Brenner, I. Spector, and E.D. Korn. 1987. Inhibition of actin polymerization by latrunculin A. FEBS Lett. 213:316–318.

12. Cullen, P.J., and F. Steinberg. 2018. To degrade or not to degrade: mechanisms and significance of endocytic recycling. Nat Rev Mol Cell Biol. 19:679–696.

13. de Beco, S., C. Gueudry, F. Amblard, and S. Coscoy. 2009. Endocytosis is required for E- cadherin redistribution at mature adherens junctions. Proc Natl Acad Sci U S A. 106:7010–7015.

14. Desclozeaux, M., J. Venturato, F.G. Wylie, J.G. Kay, S.R. Joseph, H.T. Le, and J.L. Stow. 2008. Active Rab11 and functional recycling endosome are required for E-cadherin trafficking and lumen formation during epithelial morphogenesis. Am J Physiol Cell Physiol. 295:C545–556.

15. Dongre, A., and R.A. Weinberg. 2019. New insights into the mechanisms of epithelial- mesenchymal transition and implications for cancer. Nat Rev Mol Cell Biol. 20:69–84.

16. Dunst, S., T. Kazimiers, F. von Zadow, H. Jambor, A. Sagner, B. Brankatschk, A. Mahmoud, S. Spannl, P. Tomancak, S. Eaton, and M. Brankatschk. 2015. Endogenously tagged rab proteins: a resource to study membrane trafficking in *Drosophila*. Dev Cell. 33:351–365.

17. Franz, A., and V. Riechmann. 2010. Stepwise polarisation of the *Drosophila* follicular epithelium. Developmental biology. 338:136–1347.

18. Grindstaff, K.K., C. Yeaman, N. Anandasabapathy, S.C. Hsu, E. Rodriguez-Boulan, R.H. Scheller, and W.J. Nelson. 1998. Sec6/8 complex is recruited to cell-cell contacts and specifies transport vesicle delivery to the basal-lateral membrane in epithelial cells. Cell. 93:731–740.

19. Gutzeit, H.O. 1990. The microfilament pattern in the somatic follicle cells of mid-vitellogenic ovarian follicles of *Drosophila*. Eur J Cell Biol. 53:349–356.

20. Hanson, B.J., and W. Hong. 2003. Evidence for a role of SNX16 in regulating traffic between the early and later endosomal compartments. J Biol Chem. 278:34617–34630.

21. Harris, T.J., and U. Tepass. 2010. Adherens junctions: from molecules to morphogenesis. Nature reviews. Molecular cell biology. 11:502–514.

22. Hill, E., J. van Der Kaay, C.P. Downes, and E. Smythe. 2001. The role of dynamin and its binding partners in coated pit invagination and scission. J Cell Biol. 152:309–323.

23. Horne-Badovinac, S., and D. Bilder. 2005. Mass transit: epithelial morphogenesis in the *Drosophila* egg chamber. Dev Dyn. 232:559–574.

24. Huang, J., W. Zhou, W. Dong, A.M. Watson, and Y. Hong. 2009. From the Cover: Directed, efficient, and versatile modifications of the Drosophila genome by genomic engineering. Proc Natl Acad Sci U S A. 106:8284–8289.

25. Jin, Y., A. Sultana, P. Gandhi, E. Franklin, S. Hamamoto, A.R. Khan, M. Munson, R. Schekman, and L.S. Weisman. 2011. Myosin V transports secretory vesicles via a Rab GTPase cascade and interaction with the exocyst complex. Dev Cell. 21:1156–1170.

26. Kametani, Y., and M. Takeichi. 2007. Basal-to-apical cadherin flow at cell junctions. Nat Cell Biol. 9:92–98.

27. Klöpper, T.H., N. Kienle, D. Fasshauer, and S. Munro. 2012. Untangling the evolution of Rab G proteins: implications of a comprehensive genomic analysis. BMC Biol. 10:71.

28. Krauss, J., S. López de Quinto, C. Nüsslein-Volhard, and A. Ephrussi. 2009. Myosin-V regulates oskar mRNA localization in the *Drosophila* oocyte. Curr Biol. 19:1058–1063.

29. Laiouar, S., N. Berns, A. Brech, and V. Riechmann. 2020. RabX1 Organizes a Late Endosomal Compartment that Forms Tubular Connections to Lysosomes Consistent with a “Kiss and Run” Mechanism. Curr Biol. 30:1177–1188.e1175.

30. Langevin, J., M.J. Morgan, J.B. Sibarita, S. Aresta, M. Murthy, T. Schwarz, J. Camonis, and Y. Bellaiche. 2005. *Drosophila* exocyst components Sec5, Sec6, and Sec15 regulate DE-Cadherin trafficking from recycling endosomes to the plasma membrane. Developmental cell. 9:365–376.

31. Le Droguen, P.M., S. Claret, A. Guichet, and V. Brodu. 2015. Microtubule-dependent apical restriction of recycling endosomes sustains adherens junctions during morphogenesis of the *Drosophila* tracheal system. Development. 142:363–374.

32. Le, T.L., A.S. Yap, and J.L. Stow. 1999. Recycling of E-cadherin: a potential mechanism for regulating cadherin dynamics. J Cell Biol. 146:219–232.

33. Lock, J.G., and J.L. Stow. 2005. Rab11 in recycling endosomes regulates the sorting and basolateral transport of E-cadherin. Mol Biol Cell. 16:1744–1755.

34. Margolis, J., and A. Spradling. 1995. Identification and behavior of epithelial stem cells in the *Drosophila* ovary. Development. 121:3797–3807.

35. Michel, M., I. Raabe, A.P. Kupinski, R. Pérez-Palencia, and C. Bökel. 2011. Local BMP receptor activation at adherens junctions in the *Drosophila* germline stem cell niche. Nat Commun. 2:415.

36. Oda, H., T. Uemura, Y. Harada, Y. Iwai, and M. Takeichi. 1994. A *Drosophila* homolog of cadherin associated with armadillo and essential for embryonic cell-cell adhesion. Dev Biol. 165:716–726.

37. Pirraglia, C., J. Walters, and M.M. Myat. 2010. Pak1 control of E-cadherin endocytosis regulates salivary gland lumen size and shape. Development. 137:4177–4189.

38. Pocha, S.M., T. Wassmer, C. Niehage, B. Hoflack, and E. Knust. 2011. Retromer controls epithelial cell polarity by trafficking the apical determinant Crumbs. Curr Biol. 21:1111–1117.

39. Prasad, M., A.C. Jang, M. Starz-Gaiano, M. Melani, and D.J. Montell. 2007. A protocol for culturing *Drosophila* melanogaster stage 9 egg chambers for live imaging. Nature protocols. 2:2467–2473.

40. Prigent, M., T. Dubois, G. Raposo, V. Derrien, D. Tenza, C. Rosse, J. Camonis, and P. Chavrier. 2003. ARF6 controls post-endocytic recycling through its downstream exocyst complex effector. J Cell Biol. 163:1111–1121.

41. Riedel, F., A. Galindo, N. Muschalik, and S. Munro. 2018. The two TRAPP complexes of metazoans have distinct roles and act on different Rab GTPases. J Cell Biol. 217:601–617.

42. Riedel, F., A.K. Gillingham, C. Rosa-Ferreira, A. Galindo, and S. Munro. 2016. An antibody toolkit for the study of membrane traffic in *Drosophila melanogaster*. Biol Open. 5:987–992.

43. Rodal, A.A., A.D. Blunk, Y. Akbergenova, R.A. Jorquera, L.K. Buhl, and J.T. Littleton. 2011. A presynaptic endosomal trafficking pathway controls synaptic growth signaling. J Cell Biol. 193:201–217.

44. Roeth, J.F., J.K. Sawyer, D.A. Wilner, and M. Peifer. 2009. Rab11 helps maintain apical crumbs and adherens junctions in the *Drosophila* embryonic ectoderm. PLoS One. 4:e7634.

45. Rojas, R., T. van Vlijmen, G.A. Mardones, Y. Prabhu, A.L. Rojas, S. Mohammed, A.J. Heck, G. Raposo, P. van der Sluijs, and J.S. Bonifacino. 2008. Regulation of retromer recruitment to endosomes by sequential action of Rab5 and Rab7. J Cell Biol. 183:513–526.

46. Seaman, M.N., M.E. Harbour, D. Tattersall, E. Read, and N. Bright. 2009. Membrane recruitment of the cargo-selective retromer subcomplex is catalysed by the small GTPase Rab7 and inhibited by the Rab-GAP TBC1D5. J Cell Sci. 122:2371–2382.

47. Simonetti, B., and P.J. Cullen. 2019. Actin-dependent endosomal receptor recycling. Curr Opin Cell Biol. 56:22–33.

48. Skora, A.D., and A.C. Spradling. 2010. Epigenetic stability increases extensively during *Drosophila* follicle stem cell differentiation. Proc Natl Acad Sci U S A. 107:7389–7394.

49. Solinger, J.A., H.O. Rashid, C. Prescianotto-Baschong, and A. Spang. 2020. FERARI is required for Rab11-dependent endocytic recycling. Nat Cell Biol. 22:213–224.

50. Solis, G.P., N. Hülsbusch, Y. Radon, V.L. Katanaev, H. Plattner, and C.A. Stuermer. 2013. Reggies/flotillins interact with Rab11a and SNX4 at the tubulovesicular recycling compartment and function in transferrin receptor and E-cadherin trafficking. Mol Biol Cell. 24:2689–2702.

51. Spector, I., N.R. Shochet, Y. Kashman, and A. Groweiss. 1983. Latrunculins: novel marine toxins that disrupt microfilament organization in cultured cells. Science. 219:493–495.

52. Takeichi, M. 2014. Dynamic contacts: rearranging adherens junctions to drive epithelial remodelling. Nat Rev Mol Cell Biol. 15:397–410.

53. Tepass, U., C. Theres, and E. Knust. 1990. crumbs encodes an EGF-like protein expressed on apical membranes of Drosophila epithelial cells and required for organization of epithelia. Cell. 61:787–799.

54. Thiery, J.P., H. Acloque, R.Y. Huang, and M.A. Nieto. 2009. Epithelial-mesenchymal transitions in development and disease. Cell. 139:871–890.

55. Wang, S., Z. Zhao, and A.A. Rodal. 2019. Higher-order assembly of Sorting Nexin 16 controls tubulation and distribution of neuronal endosomes. J Cell Biol. 218:2600–2618.

56. Wang, Y., and V. Riechmann. 2007. The role of the actomyosin cytoskeleton in coordination of tissue growth during *Drosophila* oogenesis. Curr Biol. 17:1349–1355.

57. Weeratunga, S., B. Paul, and B.M. Collins. 2020. Recognising the signals for endosomal trafficking. Curr Opin Cell Biol. 65:17–27.

58. Woichansky, I., C.A. Beretta, N. Berns, and V. Riechmann. 2016. Three mechanisms control E-cadherin localization to the zonula adherens. Nat Commun. 7:10834.

59. Wu, S., S.Q. Mehta, F. Pichaud, H.J. Bellen, and F.A. Quiocho. 2005. Sec15 interacts with Rab11 via a novel domain and affects Rab11 localization in vivo. Nat Struct Mol Biol. 12:879–885.

60. Wu, X., B. Bowers, K. Rao, Q. Wei, and J.A. Hammer, 3rd. 1998. Visualization of melanosome dynamics within wild-type and dilute melanocytes suggests a paradigm for myosin V function In vivo. J Cell Biol. 143:1899–18918.

61. Wucherpfennig, T., M. Wilsch-Brauninger, and M. Gonzalez-Gaitan. 2003. Role of *Drosophila* Rab5 during endosomal trafficking at the synapse and evoked neurotransmitter release. J Cell Biol. 161:609–624.

62. Xu, J., L. Lan, N. Bogard, C. Mattione, and R.S. Cohen. 2011. Rab11 is required for epithelial cell viability, terminal differentiation, and suppression of tumor-like growth in the *Drosophila* egg chamber. PloS one. 6:e20180.

63. Xu, J., L. Zhang, Y. Ye, Y. Shan, C. Wan, J. Wang, D. Pei, X. Shu, and J. Liu. 2017. SNX16 Regulates the Recycling of E-Cadherin through a Unique Mechanism of Coordinated Membrane and Cargo Binding. Structure. 25:1251–1263.e1255.

64. Yeaman, C., K.K. Grindstaff, and W.J. Nelson. 2004. Mechanism of recruiting Sec6/8 (exocyst) complex to the apical junctional complex during polarization of epithelial cells. J Cell Sci. 117:559–570.

65. Zhang, J., K.L. Schulze, P.R. Hiesinger, K. Suyama, S. Wang, M. Fish, M. Acar, R.A. Hoskins, H.J. Bellen, and M.P. Scott. 2007. Thirty-one flavors of *Drosophila* rab proteins. Genetics. 176:1307–1322.

66. Zhang, X.M., S. Ellis, A. Sriratana, C.A. Mitchell, and T. Rowe. 2004. Sec15 is an effector for the Rab11 GTPase in mammalian cells. J Biol Chem. 279:43027–43034.

67. Zobel, T., K. Brinkmann, N. Koch, K. Schneider, E. Seemann, A. Fleige, B. Qualmann, M.M. Kessels, and S. Bogdan. 2015. Cooperative functions of the two F-BAR proteins Cip4 and Nostrin in the regulation of E-cadherin in epithelial morphogenesis. J Cell Sci. 128:1453.

